# Increased perceived effort during contralateral thermal heat pain is not explained by increased intracortical and corticospinal inhibition

**DOI:** 10.64898/2026.06.30.735616

**Authors:** Ilaria Monti, Maxime Bergevin, Madhumitha Murugavel Sangeetha, Thomas Mangin, Jason Neva, Mathieu Roy, Pierre Rainville, Benjamin Pageaux

## Abstract

**Background:** Pain influences motor function and has been proposed to reduce corticospinal and intracortical excitability. At the same time, performance can be maintained during pain, at the cost of increased perceived effort, a centrally generated signal reflecting resource engagement. Here, we tested whether contralateral thermal heat pain-related changes in corticospinal and intracortical excitability contribute to increased effort perception.

**Methods:** In this preregistered transcranial magnetic stimulation (TMS) study, twenty-one healthy participants received single and paired pulse TMS at rest and during submaximal isometric right wrist flexions performed at 20% maximal peak force. Trials were conducted under a control condition or during contralateral thermal stimulation (painful or non-painful warm) applied to the left forearm. After each contraction, participants rated the intensity of their perceived effort. Corticospinal and intracortical excitability of the right wrist flexor was assessed at rest and during submaximal contractions.

**Results:** Contralateral heat pain significantly increased perceived effort compared with the control and warm conditions. Contralateral heat pain did not reduce corticospinal or intracortical excitability. Conversely, contralateral heat pain increased corticospinal excitability, reflected primarily in decreased cortical silent period duration. Perceived effort was associated with the subjective experience of pain rather than with TMS-derived variables.

**Conclusions:** These findings suggest that increased effort during contralateral heat pain cannot be attributed to inhibition of the primary motor cortex or the corticospinal pathway. The higher perceived effort in the presence of contralateral heat pain likely reflects the cognitive cost of pain rather than alterations in the transmission of the motor command.

## Introduction

Pain attracts attention (Legrain et al., 2009) and influences the motor system via spinothalamic projections to motor areas, including the supplementary motor area (SMA) and midcingulate cortex (MCC) (Dum et al., 2009). By altering cognitive and motor processes, pain increases perceived task difficulty (Cancela et al., 2023). Motivational intensity theory proposes that effort scales with task difficulty (Richter et al., 2016). Thus, maintaining task performance during submaximal tasks under pain requires increased effort.

Supporting this possibility, recent work showed that visuomotor performance can be maintained during contralateral heat pain, with pain-related costs reflected as increased perceived effort (Mangin et al., 2026). This increase was observed across visuomotor and cognitive tasks, suggesting that compensatory effort during pain may generalize across domains. Subsequent neuroimaging work replicated this behavioral effect and linked pain-related increases in perceived effort during visuomotor performance to altered SMA and MCC activity (Monti et al., 2026), regions implicated in central motor command generation (Amador & Fried, 2004; Gillies et al., 2019) and cognitive control (Botvinick et al., 2001).

Effort reflects the voluntary engagement of physical and cognitive resources to achieve a goal (Mangin & Pageaux, 2026; Preston & Wegner, 2009). Effort perception is the subjective experience of engaged resources, informing about task difficulty and intensity (Mangin & Pageaux, 2026). Converging evidence indicates that effort perception arises from the brain processing of corollary discharge signals associated with motor commands (Bergevin et al., 2023; De Morree & Marcora, 2015). Within this framework, increased perceived effort during pain may reflect altered generation or transmission of motor commands (Mangin et al., 2026). Interestingly, while previous work observed altered SMA and MCC activity during painful visuomotor task execution, suggesting altered motor command related to the cognitive control cost of pain (Monti et al., 2026), increased perceived effort may also reflect altered motor command transmission.

Transcranial magnetic stimulation (TMS) non-invasively assesses the primary motor cortex functional state and descending corticospinal pathway by evaluating intracortical and corticospinal excitability. Corticospinal excitability, indexed by motor-evoked potential (MEP) amplitude and cortical silent period (CSP) duration, reflects cortical and spinal neurons capacity to transmit motor commands (Badawy et al., 2012). Intracortical excitability, assessed with short-interval intracortical inhibition (SICI) and intracortical facilitation (ICF), reflects inhibitory and excitatory cortical circuits involved in motor command regulation (Ni & Chen, 2012). Recent meta-analyses reported inhibitory effect of experimental pain on corticospinal excitability (Chowdhury et al., 2022; Rohel et al., 2021). Whether such inhibition contributes to increased perceived effort remains unclear, as maintaining motor output under inhibition may require increasing central motor command.

This preregistered study investigated how contralateral thermal heat pain modulates corticospinal and intracortical excitability. Because corticospinal and intracortical excitability differ between rest and active conditions (Rossini et al., 1994), and effort requires active task engagement (Mangin & Pageaux, 2026), we tested pain effects at rest and during submaximal isometric wrist flexions. We also examined whether pain-related corticospinal and intracortical excitability changes were associated with increased perceived effort. We hypothesized pain reduces corticospinal and intracortical excitability and contributes to higher perceived effort.

## Materials and methods

### Participants

Twenty-one healthy, right-handed volunteers were recruited for the study (11 females and 10 males; mean age F: 27.73 ± 5.82 years, M: 27.20 ± 5.12 years; age range 18–40 years; mean weight F: 54.31 ± 7.43 kg, M: 71.50 ± 4.90 kg; mean height F: 162.36 ± 6.15 cm, M: 172.97 ± 5.94 cm). A sensitivity analysis for a repeated measure ANOVA with options “as in G*Power 3.0”, alpha risk =.05, sample size of 20, number of measurements = 3 and correlation among repeated measures =.5 indicated that we have 90% chance to observe an effect size of f = 0.338 or higher (which is equal to *ƞ_p_²* = 0.1, medium to large effect size).

Exclusion criteria included any history of pain-related, neurological or psychiatric disorders, serious medical conditions (e.g., cancer or cardiovascular disease), functional musculoskeletal impairments, and current use of psychotropic or analgesic medications. All participants provided written informed consent and received a compensation of 120 CAD for their time and commitment to the study. The study protocol was approved by the Research Ethics Committee of the “Institut Universitaire de Gériatrie de Montréal”.

### Experimental design

This study was preregistered on the Open Science Framework prior to data analysis (registration link: https://osf.io/eq2zv).

Participants visited the laboratory three times: Visit 1: calibration session, Visit 2: rest TMS session, Visit 3: active TMS session. The order of visits 2 and 3 was randomized and counterbalanced across participants.

The first visit served as a calibration and familiarization session, during which each participant underwent sensory calibration of thermal stimulations, maximal voluntary contraction (MVC) peak torque measurement, and familiarization with isometric wrist flexions. These procedures are described in detail below.

During rest and active TMS sessions, electromyographic signals were recorded from right flexor carpi radialis (FCR) and extensor carpi radialis (ECR) muscles either at rest or during isometric wrist flexions at 20% of maximal peak torque. In both sessions, TMS was applied in a control condition (no thermal stimulation) and during non-painful (warm) or painful thermal stimulation applied to the left forearm.

### Experimental setup

Participants were seated on an adjustable chair facing a computer screen displaying the target force feedback line to be matched during the active session. The right forearm was positioned in a dynamometer, and the chair was individually adjusted to maintain a consistent elbow angle at ∼120° across sessions. The left forearm rested comfortably on the desk with the palm facing upward. Participants maintained an upright posture and wore a neck collar to stabilize the head during TMS procedures. **Fig.1a** provides a schematic illustration of the experimental setup.

### Rest and Active TMS sessions

Rest and active TMS sessions followed the same overall structure (**Fig. 1b**), with the key difference being that during the *rest session*, TMS measurements were recorded with the right forearm positioned in a wrist dynamometer while participants remained at rest, without performing any voluntary contraction. During the *active session*, measurements were obtained while participants executed isometric right wrist flexions at 20% of their maximal voluntary contraction (MVC) peak torque. During both sessions, TMS-evoked responses were recorded from the right FCR and ECR.

**Figure 1:**
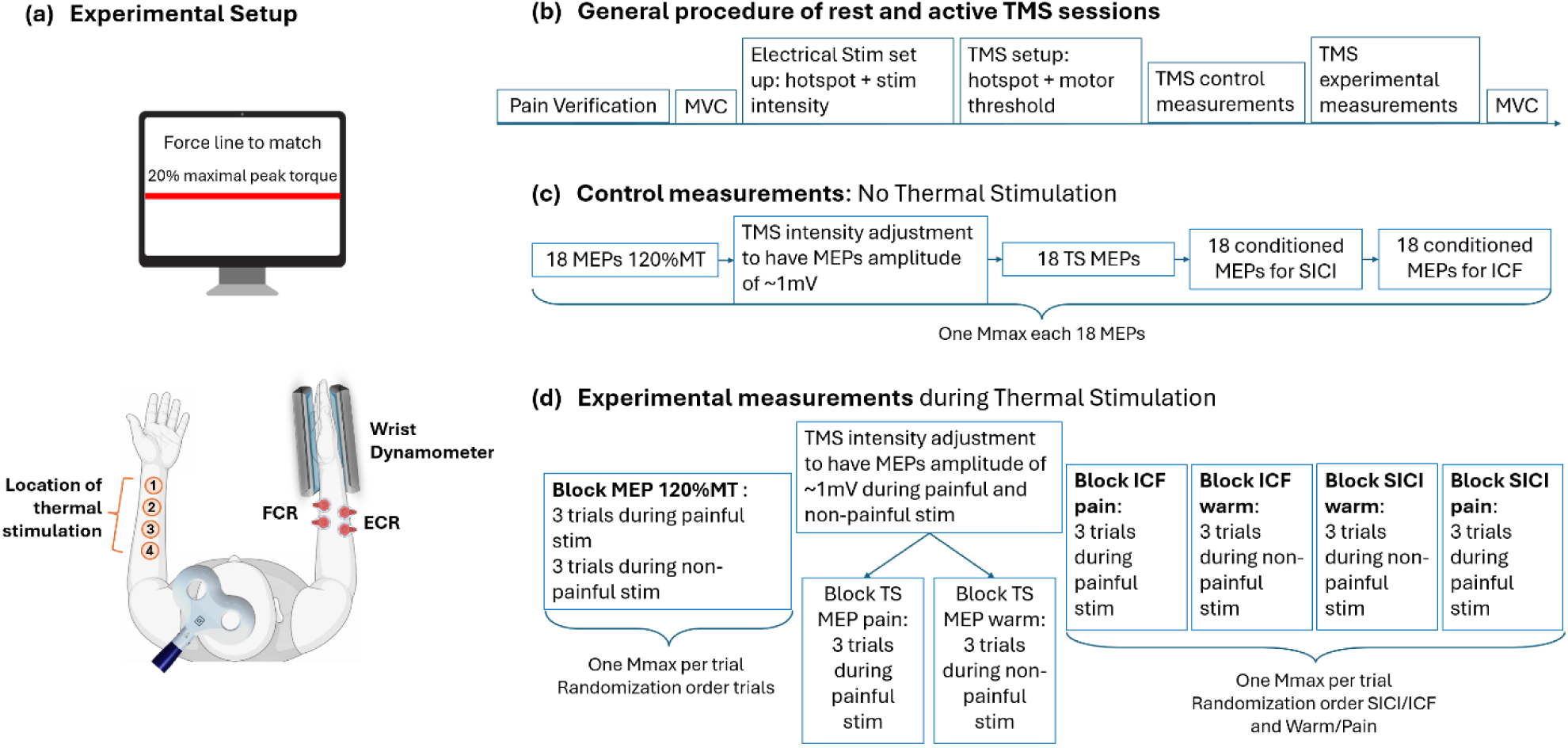
Experimental design. *Panel (a) Experimental setup.* Participants were seated facing a computer screen displaying the force line to be matched during active trials. The right forearm was positioned in a wrist dynamometer with the thumb upward and fingers extended in the sagittal plane. Four electrodes recording electromyographic (EMG) activity from flexor carpi radialis (FCR) and extensor carpi radialis (ECR) muscles. The left forearm rested on the desk, palm upward, with four marked sites for thermal stimulation. Participants maintained an upright posture and wore a neck collar for head stabilization. The transcranial magnetic stimulation (TMS) coil was positioned tangentially over the left primary motor cortex. *Panel (b) General procedure of rest and active TMS sessions.* Pain verification: Participants were exposed to the target painful and non-painful temperature to ensure consistent perception of high pain (70/100) across sessions; pain temperature was adjusted if necessary. Maximal voluntary contraction (MVC): Participants performed three maximal isometric wrist flexions; the highest peak torque was used to set wrist flexions at 20% MVC during the active session. Electrical stimulation setup: The optimal radial nerve hotspot on the right arm was identified to elicit the largest M-wave and to set the stimulation intensity. TMS setup: The optimal left primary motor cortex hotspot was determined to measure resting or active motor threshold. TMS control measurements: All TMS measurements were recorded in the absence of any thermal stimulation. TMS experimental measurements: All TMS measurements were recorded during non-painful and painful thermal stimulations. *Panel (c) Control measurements.* Order of TMS measurements without thermal stimulation. One maximal compound muscle action potential (Mmax) was recorded after each 18 MEPs. *Panel (d) Experimental measurements.* Order of TMS measurements during painful or non-painful thermal stimulation. One Mmax was recorded at the end of each trial, see also Fig.2. For both rest and active TMS sessions, control measurements preceded experimental measurements. For experimental measurements, the order of trials in the MEP 120% motor threshold (MT) block was randomized; the short intracortical inhibition (SICI) and intracortical facilitation (ICF) protocols were presented in randomized order, and within each protocol, the order of warm and pain blocks was counterbalanced across participants.

For both sessions and for each TMS protocol, control measures were obtained by delivering 18 TMS pulses in the absence of thermal stimulation (**Fig. 1c**). Experimental measurements comprised 24 trials with individually calibrated painful or non-painful thermal stimulation, during which TMS pulses were delivered (**Fig. 1d**). Experimental trials were organized into blocks corresponding to different TMS protocols. Block MEP consisted of six trials to record MEP at 120% resting or active motor threshold during painful and non-painful thermal stimulations. Test stimulus (TS) MEP pain block and TS MEP warm block consisted of three trials each to record TS MEP during painful and non-painful thermal stimulations, respectively. For the ICF and SICI protocols, separate pain and warm blocks (three trials each) were used to record conditioned MEPs during painful and non-painful stimulation.

For MEP block, the order of thermal stimulation was randomized across trials. For SICI and ICF, the order of blocks was randomized across participants (half started with SICI, half with ICF). Within SICI and ICF blocks, the order of thermal stimulation was also randomized, such that half of the participants experienced pain first and half experienced warm first.

### Experimental Trials

Experimental trials consisted of two types of thermal stimulation conditions: (i) warm condition, during which non-painful thermal stimulation was delivered; (ii) pain condition, during which painful thermal stimulation was delivered. Each trial consisted of 30 s of thermal stimulation; participants either remained at rest (**Fig. 2a**, rest session) or performed 30 s of isometric wrist flexion at 20% MVC peak torque (**Fig. 2b**, active session). At the beginning of each trial, an auditory cue (beep) signaled the onset of thermal stimulation and either the wrist flexion (active trial) or the rest period (rest trial).

**Figure 2:**
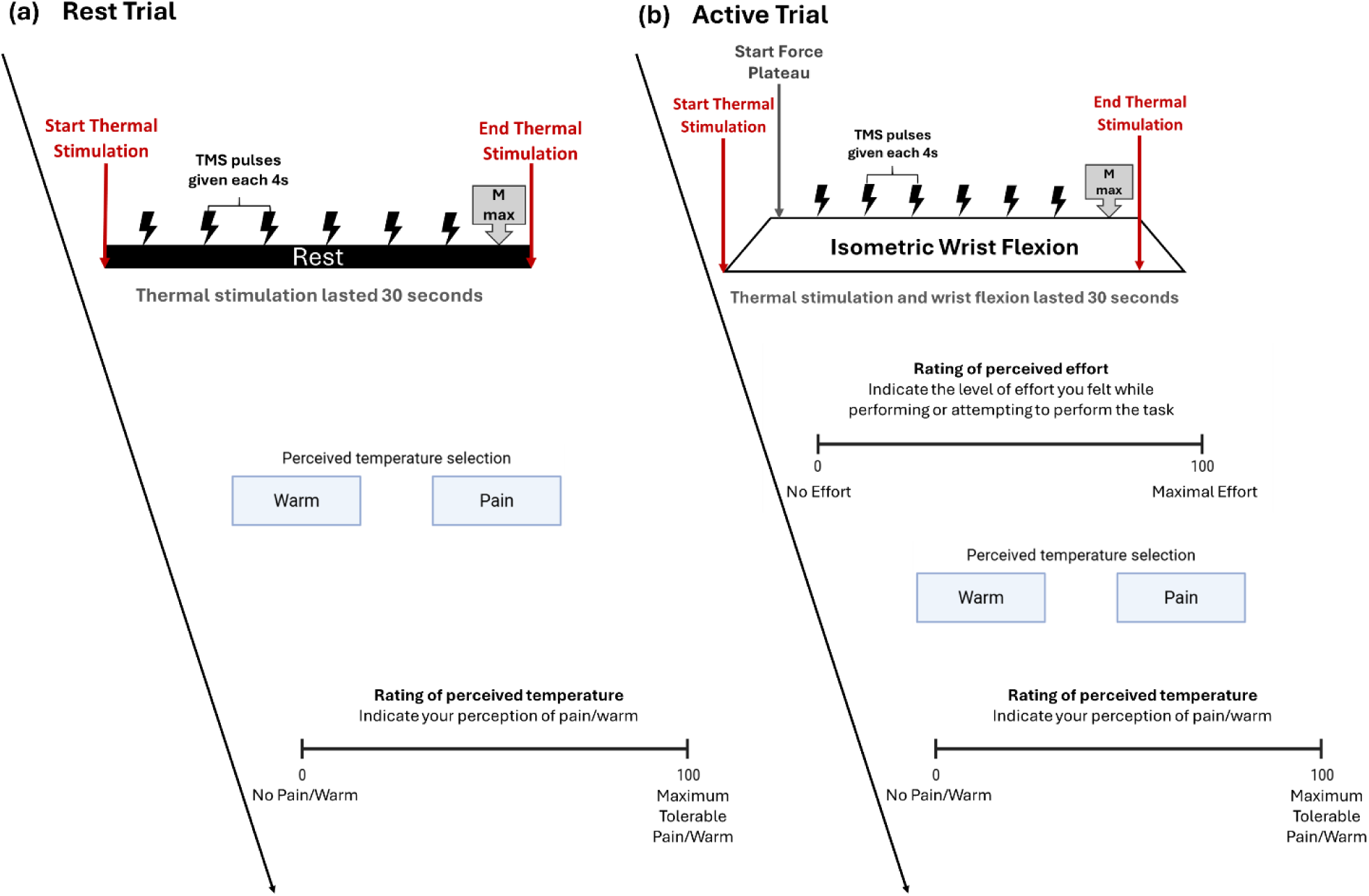
Example of trials. *Panel (a) Rest trial.* Each trial consisted of 30 s of thermal stimulation during which six transcranial magnetic stimulation (TMS) pulses were delivered at 4-s intervals. Approximately 28 s after stimulation onset, an electrical stimulus elicited a maximal compound muscle action potential (Mmax). Following the offset of thermal stimulation, participants indicated whether it was warm or painful and rated its intensity on a 0–100 visual analog scale. *Panel (b) Active trial.* An auditory cue marked the onset of thermal stimulation and wrist flexion. Once participants maintained 20% of their maximal torque (force plateau), six TMS pulses were delivered at 4-s intervals. At ∼28 s, an electrical stimulus elicited an Mmax. Both thermal stimulation and wrist flexion lasted 30 s, the offset was indicated by a second auditory cue. Participants then rated perceived effort on a 0–100 visual analog scale, indicated whether the thermal stimulation was warm or painful by choosing between two buttons, and rated temperature intensity on a 0–100 visual analog scale.

During active trials, TMS pulses were delivered at the force plateau, no TMS pulses were delivered if force output fell outside the target range as visually verified by the experimenter delivering the TMS pulse. In both active and rest trials, a peripheral nerve electrical stimulation was delivered approximately 28s after trial onset to elicit a maximal compound muscle action potential (Mmax). Each active trial was followed by a 1 min recovery period to limit the development of neuromuscular fatigue.

Rest trials were followed by a rating period to record temperature perception. Active trials were followed by ratings of perceived effort and temperature.

Perception of effort was defined as the effort experienced to match the force feedback line as steadily and precisely as possible (Mangin et al., 2026; Monti et al., 2026). Participants rated the intensity of their perceived effort using a visual analog scale (VAS; 0–100). Thermal stimulation ratings required participants to first indicate whether the stimulus was perceived as painful or non-painful by selecting one of two buttons (“pain” or “warm”) and then rate the perceived intensity of pain or warmth using a VAS (0–100). Between trials, the thermal stimulator was moved to one of four different sites on the left forearm to prevent local sensitization.

### Sensory calibration and thermal stimulations

Thermal stimuli were delivered using a 3×3cm contact thermode applied to 4 skin sites on the volar surface of the left non-dominant forearm (TCS unit and T11 probe, QST.Lab, Strasbourg, France). Stimuli were delivered between 42°C (slightly warm) and 50°C (max) with a plateau duration of 30s. The maximum temperature of 50°C typically produces strong but tolerable pain and poses no risk of skin damage.

Painful thermal stimuli were individually calibrated by first estimating a temperature of pain tolerance and then applying thermal stimuli between 42°C and the temperature of pain tolerance. Then, the target pain temperature was chosen to produce a pain intensity rated at ∼70/100 on a pain visual analog scale (VAS, from 0 = “no pain at all” to 100 = “maximum tolerable pain”. For all participants, the non-painful temperature was set to 41.9°C. This temperature avoids the activation of nociceptors and generally produces a non-painful warm sensation (Mangin et al., 2026; Monti et al., 2026).

Prior to the visits 2 and 3 (i.e., TMS visits), participants were re-exposed to the target painful and non-painful temperature and rated their perceptions of warmth and pain using the VAS. This assessment ensured that the perception of pain and warmth was consistent between sessions. If necessary, pain temperature was adjusted prior to the onset of TMS sessions to maintain a high pain rating of approximately 70/100 on the VAS.

At the end of each trial, the experimenter moved the thermal stimulator to a different arm spot (4 different arm spots were used) to prevent a possible sensitization of the stimulated area.

### Maximal Voluntary Contraction

The maximal voluntary contraction (MVC) peak torque was collected with a custom-made wrist dynamometer by S2P (Science to Practice, Ltd, Slovenia) connected to a PowerLab system (PL3516 PowerLab 16/35, AD Instrument, Colorado Springs, CO, USA). The torque signal was recorded at a sample frequency of 2 kHz and stored for offline analysis.

The right wrist was placed in the dynamometer in a neutral anatomical position. The wrist was stabilized at the level of hand palm, wrist and elbow with vertical beams covered with foams, to avoid movements and ensure the same position across trials and visits. The thumb pointed upward, and the fingers were extended in a neutral position in the sagittal plane.

At the beginning and the end of the rest and active TMS sessions, participants performed three maximal isometric wrist flexions. Changes in MVC peak torque were used to evaluate neuromuscular fatigue, allowing verification that TMS stimulation and/or repeated isometric flexions did not induce muscle fatigue that could confound the results.

During the active TMS session, the force of the isometric wrist flexions was set at the 20% MVC peak torque, based on the highest MVC peak torque obtained at the beginning of the session.

### Submaximal isometric wrist flexions

Voluntary submaximal isometric wrist flexions were performed at 20% of individual MVC peak torque measured at the beginning of each visit. The feedback of the force produced during the isometric wrist flexion was displayed on a screen, allowing participants to adjust their force to match the target torque. Each isometric flexion was continuously performed during 30s with the right wrist in the same position adopted during the MVC.

Each 30 s isometric flexion was followed by a pause to record perception of effort and perception of pain/warm as well as to minimize the development of muscle fatigue. Wrist flexions at 20%MVC for 30 s should not induce muscular fatigue, which seems to appear only after 50/70s of isometric wrist flexion at 25%MVC (Forman et al., 2020), or after 1-min of handgrip contraction at 25% MVC (Byström & Kilbom, 1990).

The force produced during each flexion was collected with the custom-made wrist dynamometer by S2P (Science to Practice, Ltd, Slovenia) connected to a PowerLab system (PL3516 PowerLab 16/35, AD Instruments, Colorado Springs, CO, USA). The torque signal was recorded at a sample frequency of 2 kHz and stored for offline analysis. Visual feedback of the force produced during the contraction was provided on the screen using a colored bar. Participants were requested to match as precisely and steadily as possible the colored force target bar during 30 s.

### Electromyographic (EMG) recording

For all TMS measurements, surface EMG was recorded from the right flexor carpi radialis (FCR) and the right extensor carpi radialis (ECR) muscles using bipolar Ag/AgCl electrodes of 1 cm diameter placed longitudinally along the muscle–tendon axis. Electrode pairs were positioned to the muscle belly closer to the tendinous region, to obtain the cleanest MEP and Mmax wave form. The ground (reference) electrode was placed over the lateral epicondyle of the humerus.

Surface EMG was recorded using LabChart software (LabChart 8.0). EMG signal was sampled using a PowerLab (PL3516 PowerLab, 16/35, AD Instruments, Colorado Springs, CO, USA) data acquisition system and a bioamplifier (Dual Bio Amp, AD Instruments, Colorado Springs, CO, USA) with an acquisition rate of 2 kHz, 10 to 500 Hz bandpass and 60Hz notch filter.

### Transcranial Magnetic Stimulation

We administered single-pulse and paired-pulse TMS over the left primary motor cortex (M1) using a Magstim BiStim 200^2^ stimulator (Magstim Co., UK) connected to a figure-of-eight magnetic coil (Magstim 70 mm P/N 9790, Magstim Co., UK) and recorded TMS-induced MEPs from the contralateral right flexor carpi radialis (FCR) and extensor carpi radialis (ECR) using PowerLab electromyography (EMG) data acquisition system (AD Instruments, CO, USA).

The TMS coil was held tangentially to the skull, with the handle pointing backward and laterally at approximately 45° to the midline, resulting in a posterior-anterior direction of current flow in the brain. The optimal coil position over left M1 was determined for each participant and defined as the point on the scalp where stimulation consistently evokes the largest MEPs in the right FCR. The optimal coil position over left M1 was marked on a template brain present in Brainsight neuronavigation software (Rogue Research Inc., Montreal, QC, Canada), used to monitor coil position and ensure consistent stimulation across trials for each participant.

#### Single-pulse TMS

Single-pulse TMS was delivered during the rest session to collect MEPs and during the active session to collect MEPs and CSP both from FCR and ECR muscles. For MEPs recorded at rest, TMS intensity was set at 120% of the resting motor threshold (rMT), defined as the lowest stimulus intensity able to evoke MEPs with peak-to-peak amplitude of ∼50mV in the relaxed FCR muscle in 5 out of 10 consecutive stimuli (Rossini et al., 2015). For MEPs recorded during wrist flexions, TMS intensity was set at 120% of active motor threshold (aMT), defined as the lowest stimulus intensity able to induce MEPs with peak-to-peak amplitude of ∼200mV in the FCR muscle contracted at 5%MVC in 5 out of 10 consecutive stimuli (Rossini et al., 2015; Stagg et al., 2011). The intensity of stimulation for the rMT and aMT was 56 ± 10% and 51 ± 10% of the maximum stimulator output, respectively.

#### Paired-pulse TMS

Paired-pulse TMS was delivered during rest and wrist flexions to collect SICI and ICF from FCR and ECR muscles. Both SICI and ICF were measured by applying a subthreshold conditioning stimulus (CS) followed by a suprathreshold test stimulus (TS). The MEP elicited by this sequence of stimulation (CS followed by TS) is referred to as a conditioned MEP. During the rest session, the CS was delivered at 80% rMT, During the active session, the CS was delivered at 80% aMT (Ortu et al., 2008). During both TMS sessions, the TS was delivered at an intensity individually adjusted across thermal conditions to elicit an average peak-to-peak MEP amplitude of ∼1 mV across 10 stimulations (Rossini et al., 2015). For SICI the TS was applied 2ms after the CS. For ICF, the TS was applied 10ms after the CS (Kujirai et al., 1993). During the rest session, the intensity of stimulation for the TS without thermal stimulation was 78 ± 11% of the maximum stimulator output; for the TS during non-painful stimulation was 78 ± 11% of the maximum stimulator output; for the TS during painful stimulation was 80 ± 12% of the maximum stimulator output. During the active session, the intensity of stimulation for the TS without thermal stimulation was 59 ± 12% of the maximum stimulator output; for the TS during non-painful stimulation was 56 ± 10% of the maximum stimulator output; for the TS during painful stimulation was 57 ± 11% of the maximum stimulator output.

### Peripheral nerve stimulation for Mmax

Electrical stimulation was applied to a peripheral nerve to evoke maximal compound muscle action potentials (Mmax) for the FCR muscle. This Mmax was used as a normalization procedure to control for changes in EMG signal at a muscle level (Pageaux et al., 2015). The stimulation was delivered using a constant-current stimulator (Digitimer DS7R, Welwyn Garden City, UK). The stimulation site was the radial median nerve (El Bouse et al., 2013). The best hotspot of stimulation was determined according to the location that produced the largest M-wave at a given submaximal stimulus. Then, the stimulation electrodes, connected to a constant-current stimulator, were fixed at the arm by a velcro strap. The stimulation intensity was set by gradually increasing it until the M-wave amplitude reached a plateau, indicating maximal recruitment of motor units (Mmax). The stimulus intensity was set to 120% of the intensity producing the M-wave plateau (mean current, rest session 13.0 ± 4.0 mA, active session 13.7 ± 4.2 mA).

### Data processing and analysis

All TMS data were manually extracted offline using custom macro in LabChart (ADInstruments). For MEPs elicited at 120% rMT/aMT, TS MEPs, conditioned MEPs (CS–TS), and Mmax, peak-to-peak amplitudes were extracted. For all TMS measurements (120% rMT/aMT, TS MEPs, and conditioned MEPs), background EMG activity was quantified as the root mean square (RMS) of the EMG signal during the 100 ms preceding the TMS pulse. This measure was used to verify that the target muscles were relaxed during the rest TMS session and to evaluate potential effects of thermal stimulation on peripheral muscle activity that could influence MEP amplitudes. For data analysis, EMG background activity was first averaged across trials within each thermal condition, and then analyzed as raw RMS and normalized to the corresponding Mmax amplitude using the following formula:

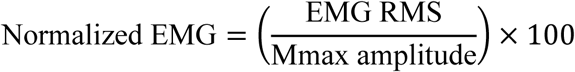

During the active TMS session, mean force output was extracted over the 100 ms preceding the TMS pulse to verify the force applied during wrist flexion was consistent at 20 ± 2%MVC peak torque. In addition, CSP duration was manually extracted by one researcher and measured in milliseconds as the interval from the TMS pulse artefact to the resumption of continuous voluntary EMG activity. Cortical silent periods were analyzed as raw duration and normalized to the MEP amplitude associated to each CSP using the following formula:

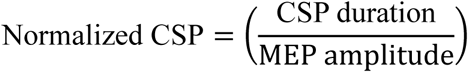

For each trial, six MEP amplitudes elicited at 120% rMT and aMT were averaged, then normalized by the subsequent Mmax amplitude included in each trial using the following formula:

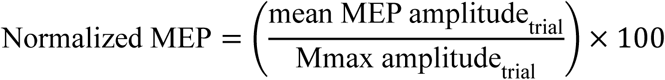

After normalization, MEP and CSP data were averaged within each thermal condition.

For TS MEPs and conditioned MEPs, peak-to-peak amplitudes were averaged directly within each thermal condition (control, warm, pain). SICI and ICF were quantified relative to the test stimulus and Mmax using the following formulas:

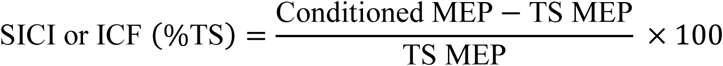

and

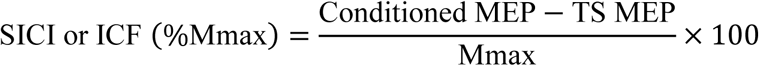

We also normalized SICI and ICF by Mmax amplitude instead of TS MEP amplitude. This approach, supported by methodological studies, allows the estimation of inhibitory processes while accounting for the proportion of spinal motoneurons contributing to MEP generation (Gueugneau et al., 2023; Lackmy & Marchand-Pauvert, 2010; Opie & Semmler, 2014).

With these two normalization procedures, negative values indicate inhibition (i.e., the conditioned MEP is smaller than the TS MEP), whereas positive values indicate facilitation (i.e., the conditioned MEP is larger than the TS MEP). Values approaching 0 reflect minimal modulation of corticospinal output relative to the TS. Accordingly, for SICI, more negative values correspond to stronger intracortical inhibition, while values closer to zero indicate reduced inhibition. For ICF, more positive values correspond to stronger intracortical facilitation, whereas values closer to zero indicate reduced facilitation.

These procedures allowed consistent quantification of corticospinal and intracortical excitability across rest and active conditions while accounting for peripheral and background EMG influences.

## Statistical analysis

The primary muscle of interest was the FCR, while the secondary muscle of interest was the ECR (antagonist involved in the contraction). An a priori approach focused on the FCR data, with the TMS and Mmax hotspot and stimulation intensity specifically optimized for this muscle. In this context, analysis of the ECR data was exploratory and reported in supplementary material.

All data were analyzed using JASP (version 0.95.4.0). Ratings of perceived effort were analyzed using a repeated-measures ANOVA with Temperature as a within-subject factor including three levels (Control, Warm, Pain). Ratings of perceived temperature were analyzed using a repeated-measures ANOVA with Temperature (Warm, Pain) and Muscle State (Rest, Active) as within-subject factors. For the pain rating, individual scores were added to a constant *k* = 100 to obtain a score > 100. The final warm and pain ratings ranged from 0 to 200, with a score below 100 indicated a warm non painful sensation, and a score above 100 indicated a painful sensation.

To evaluate the presence of muscle fatigue, paired *t*-tests were conducted separately for each session (rest and active) to compare pre-session and post-session MVC values.

To test the effect of contralateral experimental thermal heat pain on corticospinal and intracortical excitability, separate repeated-measures ANOVAs were conducted for each muscle (FCR and ECR) and muscle state (rest and active). The dependent variables were: (i) RMS of the background EMG, expressed as raw RMS or normalized RMS; (ii) MEP amplitudes elicited at 120% resting or active motor threshold, expressed as raw peak-to-peak amplitudes or normalized amplitude; (iii) CSP expressed as raw duration or normalized duration; and (iv) intracortical inhibition and facilitation indices (SICI and ICF), calculated relative to either the test stimulus or Mmax, as described in the Data analysis section; (v) mean force of the 100 ms preceding the TMS pulse during active trials; (vi) Mmax amplitude. For each ANOVA, the within-subject factor Temperature included three levels (Control, Warm, Pain). Assumptions of sphericity were tested using Mauchly’s test, and when violated, Greenhouse–Geisser–corrected results were reported. Confidence intervals (95%) for partial eta-squared (η²p) were calculated using a custom application (v1.5) developed by the author TM (https://github.com/ManginThomas/eta2p-calculator). Post hoc comparisons were corrected for multiple comparisons using the Holm–Bonferroni method.

In addition to our pre-registration, and to explore whether increases in perceived effort were associated with painful stimulation or with changes in corticospinal and intracortical excitability, repeated-measures correlations were computed. Specifically, mean perceived effort across TMS conditions was correlated with the intensity of thermal stimulation and with mean pain perception ratings. Effort perception ratings obtained during the single-pulse TMS protocol were correlated with the corresponding corticospinal measures (MEP amplitude and CSP), whereas effort perception ratings obtained during paired-pulse protocols were correlated with the corresponding intracortical measures (SICI or ICF). The repeated-measures correlations were computed using the rmcorr Shiny application (https://shinyappstore.com/a/rmcorrShiny). Corresponding *p*-values were adjusted with the Holm–Bonferroni procedure. To maintain the principle of monotonicity, if a calculated *p_adj_* was lower than the one preceding, it was set equal to the previous value, ensuring that *p_adj_*values are non-decreasing. Results were considered significant if *p_adj_*< 0.05.

## Results

### Data exclusion

One participant completed only one of the two TMS sessions. Therefore, the final dataset included data from 21 participants in the active session and 20 participants in the rest session.

At rest, MEPs contaminated by pre-stimulus EMG activity, defined as RMS EMG > 0.015mV in the 100 ms prior to the TMS pulse (Rossini et al., 2015; Shanks et al., 2023), were excluded from analysis (0.14% of all recorded FCR MEPs; 18.40% of all recorded ECR MEPs).

During the active condition, MEPs were excluded if the mean force in the 100 ms prior to the TMS pulse fell outside the target range of 18–22% MVC (1.22% of all recorded FCR and ECR MEPs).

### Manipulation check

We aimed to induce pain in order to increase the perceived effort during submaximal isometric wrist flexion.

#### Pain Perception

Painful thermal stimulation compared to non-painful stimulation augmented pain perception (main effect of temperature: *F*(1, 19) = 800.73, *p* <.001, *η²_p_* =.977 [.946,.985]) with estimated means rising from 47.4 ± 21.9 (non-painful warmth; <100) during non-painful stimulation to 164.4 ± 15.2 (painful; >100) during painful stimulation. Muscle state did not influence thermal ratings (main effect of muscle state: *F*(1, 19) = 0.17, *p* =.688, *η²_p_* =.009 [.000,.204]).

#### Effort Perception

Painful thermal stimulation compared to control condition (no thermal stimulation) and non-painful stimulation augmented perceived effort during submaximal isometric wrist flexion (**Fig. 3**; main effect of temperature: *F*(2, 40) = 3.56, *p* =.039, *η²_p_* =.151 [.000,.326]) with estimated means rising from 58.4 ± 15.3 during control condition and 58.7 ± 21.5 during non-painful stimulation, to 64.1 ± 18.7 during painful stimulation.

**Figure 3:**
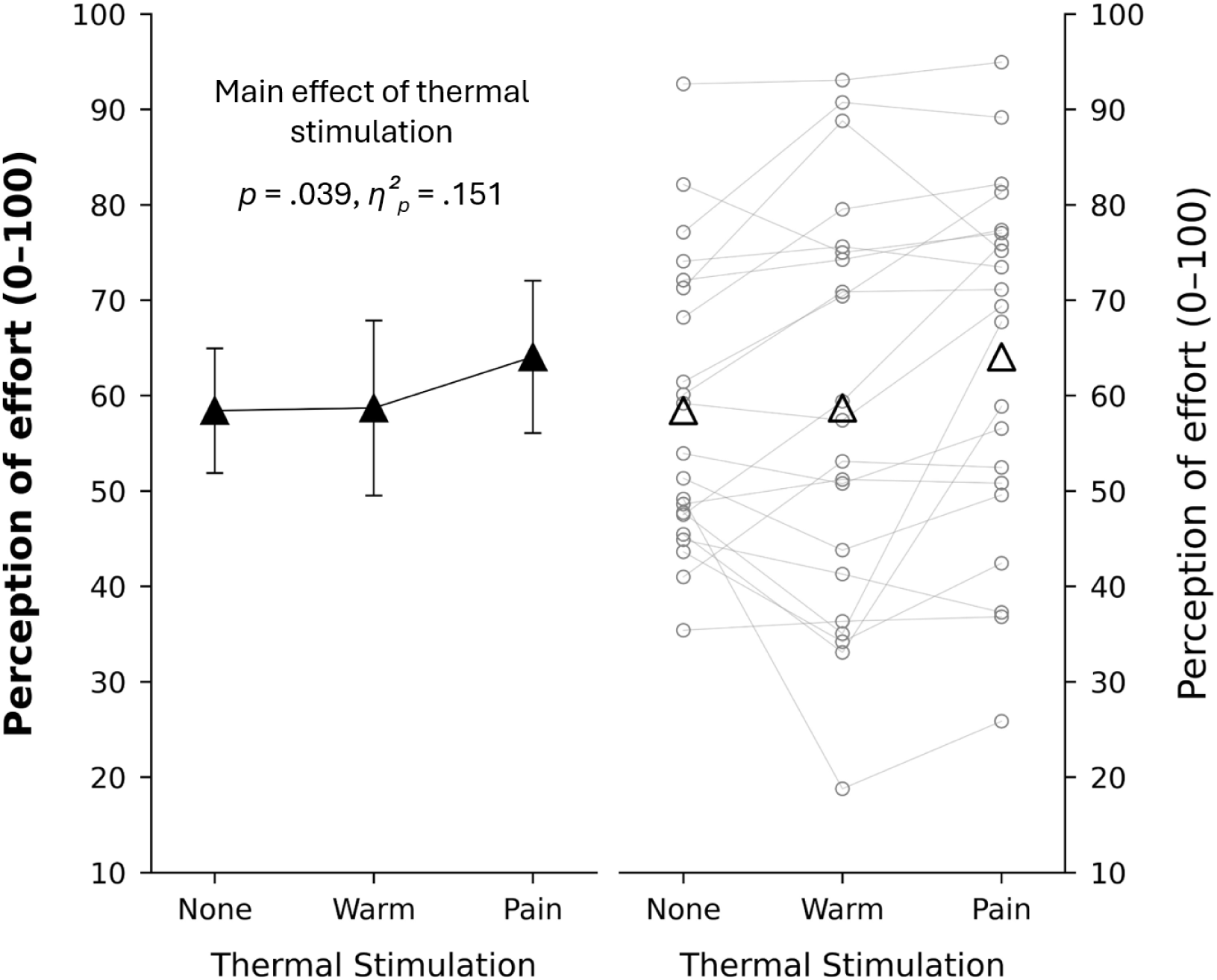
Effect of thermal stimulation on ratings of perceived effort. Ratings of perceived effort were recorded on a 0–100 visual analog scale and are shown for the three thermal stimulation conditions (none, warm, and pain). The left panel displays mean values of effort perception ratings (black triangles) with 95% confidence interval error bars. The right panel shows individual data (gray circles) and mean values (black triangles) of effort perception.

### Rest session

Maximal voluntary contraction peak force did not differ between the beginning and end of the rest session (*F*(1, 19) = 0.57, *p* =.460, *η²_p_* =.029 [.000,.260]), with estimated means going from 6.51 ± 2.69 Nm at the beginning of the session to 6.63 ± 2.93 Nm at the end of the session.

### Flexor carpi radialis

#### Mmax

During rest, mean Mmax amplitude was 4.14 ± 3.22 mV for control condition, 4.40 ± 3.27 mV for warm condition and 4.41 ± 3.28 mV for pain condition. A main effect of temperature was observed on FCR Mmax amplitude at rest (*F*(1.5, 28.70) = 5.38, *p* =.016, *η²_p_* =.221 [.007,.429]). Post hoc comparisons revealed an increase in Mmax amplitude during warm (*t*(19) = 2.63, *p* =.049, *d* = 0.080 [-0.007, 0.166]) and during pain compared to the control condition (*t*(19) = 2.43, *p* =.050, *d* = 0.084 [-0.014, 0.182]). No difference was found between the pain and warm conditions (*t*(19) = 0.21, *p* =.835, *d* = 0.004 [-0.048, 0.056]).

#### Background EMG

During rest, no main effect of temperature was found on the RMS of FCR EMG preceding TMS pulses eliciting MEPs and conditioned MEPs for SICI and ICF (all *p* >.050, all *η²_p_* <.183). Detailed statistics are presented in supplementary material 1.

#### MEP

During rest, mean raw MEP amplitude was 0.29 ± 0.12 mV for control condition, 0.28 ± 0.09 mV for warm condition and 0.34 ± 0.16 mV for pain condition. Mean normalized MEP amplitude was 13.61 ± 14.85 % for control condition, 12.69 ± 11.40 % for warm condition and 14.77 ± 14.52 % for pain condition. No main effect of temperature was found on the raw MEP amplitude (*F*(1.25, 23.67) = 2.23, *p* =.145, *η²_p_* =.105 [.000,.336]; **Fig. 4a**) nor on the normalized MEP amplitude (*F*(1.31, 24.90) = 0.821, *p* =.404, *η²_p_* =.041 [.000,.238]; **Fig. 4b**).

**Figure 4:**
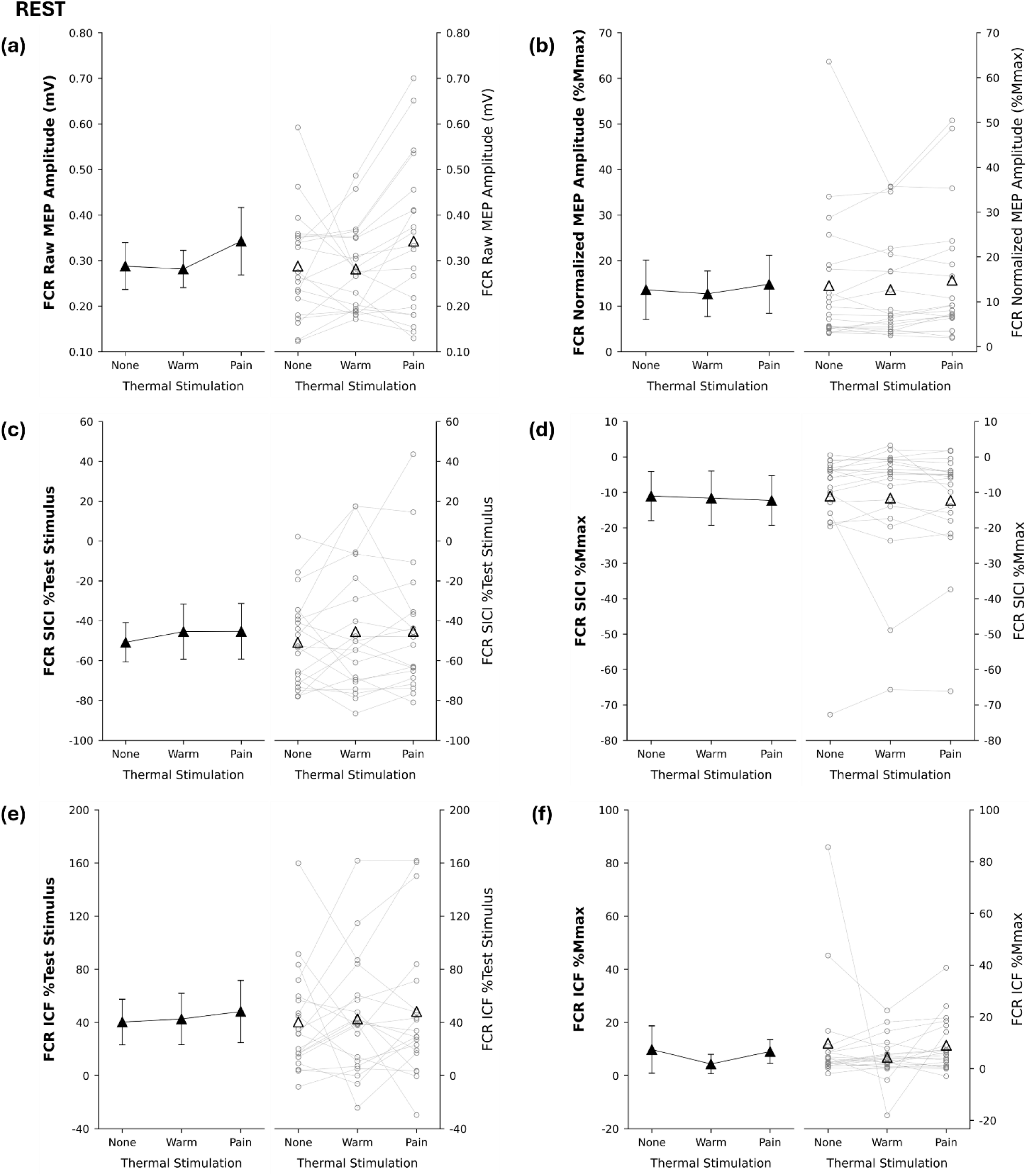
**Effect of thermal stimulation on motor evoked potential (MEP), short-interval intracortical inhibition (SICI), intracortical facilitation (ICF) recorded from the flexor carpi radialis (FCR) at rest**. In all panels, the left side shows mean values (black triangles) with 95% confidence interval error bars, while the right side shows individual data points (gray circles) along with the mean (black triangles). *Panel (a)* shows raw MEP amplitudes expressed in millivolt (mV). *Panel (b)* shows MEP amplitudes normalized by maximal compound muscle action potentials (Mmax) amplitude. *Panel (c)* shows SICI expressed in percentage of the test stimulus. *Panel (d)* shows SICI expressed in percentage Mmax amplitude. *Panel (e)* shows ICF expressed in percentage of the test stimulus. *Panel (f)* shows ICF expressed in percentage of Mmax amplitude. Repeated-measures ANOVAs revealed no main effect of thermal stimulation on MEP, SICI or ICF.

#### TS

During rest, mean test stimulus amplitude was 0.45 ± 0.20 mV for control condition, 0.47 ± 0.21 mV for warm condition and 0.56 ± 0.28 mV for pain condition. A main effect of temperature was found on test stimulus amplitude (*F*(1.86, 35.31) = 7.33, *p* =.003, *η²_p_* =.278 [.044,.462]). Post hoc comparisons revealed increased test stimulus amplitude during pain compared to control (*t*(19) = 3.39, *p* =.009, *d* = 0.484 [0.056, 0.912]) and warm conditions (*t*(19) = 2.70, *p* =.029, *d* = 0.382 [0.024, 0.788]). No significant difference was found between control and warm conditions (*t*(19) =-0.09, *p* =.378, *d* =-0.102 [-0.403, 0.199]).

#### SICI

During rest, mean SICI as percentage of test stimulus was-50.78 ± 22.48 % for control condition,-45.43 ± 31.49 for warm condition and-45.28 ± 31.77 % for pain condition. Mean SICI as percentage of Mmax was-11.04 ± 15.84 % for control condition,-11.62 ± 17.46 % for warm condition and-12.29 ± 15.95 % for pain condition. No main effect of temperature was found on SICI as percentage of test stimulus (*F*(1.91, 36.28) = 0.83, *p* =.438, *η²_p_* =.042 [.000,.186]; **Fig. 4c**), nor on SICI as percentage of the Mmax (*F*(1.34, 25.53) = 0.35, *p* =.622, *η²_p_* =.018 [.000,.183]; **Fig. 4d**).

#### ICF

During rest, mean ICF as percentage of test stimulus was 40.30 ± 39.33 % for control condition, 42.59 ± 44.11 % for warm condition and 48.22 ± 53.36 % for pain condition. Mean ICF as percentage of Mmax was 9.81 ± 20.27 % for control condition, 4.36 ± 8.32 % for warm condition and 9.06 ± 10.23 % for pain condition. No main effect of temperature was found on ICF as percentage of the test stimulus (*F*(1.79, 34.00) = 0.35, *p* =.681, *η²_p_* =.018 [.000,.140]; **Fig. 4e**), or as percentage of the Mmax (*F*(1.19, 22.52) = 1.05, *p* =.329, *η²_p_* =.053 [.000,.272]; **Fig. 4f**).

#### Extensor carpi radialis

While the hotspot and stimulation intensity were determined specifically for the FCR muscle, we also collected for exploratory analyses responses in the ECR muscle. Detailed results are presented in supplementary material 2. Briefly, at rest, we observed increased raw and normalized MEP amplitude during pain compared to warm condition (all *p* <.009, all *d* > 0.367). We also observed reduced SICI as percentage test stimulus during pain compared to warm condition (*p* =.033, *d* = 0.406). No other effect of temperature was observed.

### Active session

MVC was significantly lower at the end of the active session compared to the beginning of the same session (*F*(1, 20) = 9.75, *p* =.005, *η²_p_* =.328 [.036,.558]), with estimated means going from 6.46 ± 2.60 Nm at the beginning of the session to 5.97 ± 2.25 Nm at the end of the session.

### Flexor carpi radialis

#### Mmax

During voluntary wrist flexions, mean Mmax amplitude was 4.02 ± 3.1 mV for control condition, 4.16 ± 3.07 mV for warm condition and 4.14 ± 3.04 mV for pain condition. No main effect of temperature was found on Mmax amplitude during voluntary wrist flexions (*F*(1.24, 24.87) = 2.04, *p* =.165, *η²_p_* =.092 [.000,.315]).

#### Background EMG

Detailed statistics are presented in supplementary material 1. During voluntary wrist flexions, a main effect of temperature was observed on the RMS of FCR EMG preceding TMS pulses eliciting MEPs (*p* =.007, *η²_p_* =.262), with RMS higher during pain compared to control condition (*p* =.007, *d* = 0.215). For FCR muscle, no main effect of temperature was found on the RMS EMG preceding TMS pulses eliciting conditioned MEPs for SICI and ICF (all *p* >.428, all *η²_p_* <.036).

#### Force

Detailed statistics are presented in supplementary material 1. For FCR muscle, a main effect of temperature was observed on the mean force produced during 100 ms preceding TMS pulses eliciting MEPs (*p* =.050, *η²_p_* =.144), with mean force higher during pain (19.72 ± 0.27% MVC) compared to warm (19.67 ± 0.21% MVC) condition (*p* =.016, *d* = 0.679). No main effect of temperature was found on the mean force preceding TMS pulses eliciting conditioned MEPs for SICI and ICF (all *p* >.620, all *η²_p_* <.024).

#### MEP

During voluntary wrist flexions, mean raw MEP amplitude was 1.39 ± 0.44 mV for control condition, 1.53 ± 0.47 mV for warm condition and 1.62 ± 0.50 mV for pain condition. Mean normalized MEP amplitude was 58.45 ± 42.08 % for control condition, 56.63 ± 35.83 % for warm condition and 62.87 ± 40.42 % for pain condition. There was a main effect of temperature on raw MEP amplitude (*F*(1.28, 25.54) = 7.13, *p* =.009, *η²_p_* =.263 [.020,.482]; **Fig 5a**). Post hoc comparisons revealed that MEP amplitude was higher during pain compared to control (*t*(20) = 3.06, *p* =.019, *d* = 0.499 [0.026, 0.972]) and pain compared to warm conditions (*t*(20) = 2.83, *p* =.021, *d* = 0.193 [0.002, 0.388]), but not between control and warm conditions (*t*(20) =-2.06, *p* =.053, *d* =-0.306 [-0.714, 0.102]). These differences disappeared when MEPs amplitude was normalized to Mmax (*F*(1.30, 26.01) = 1.43, *p* =.251, *η²_p_* =.067 [.000,.275]; **Fig. 5b**).

**Figure 5:**
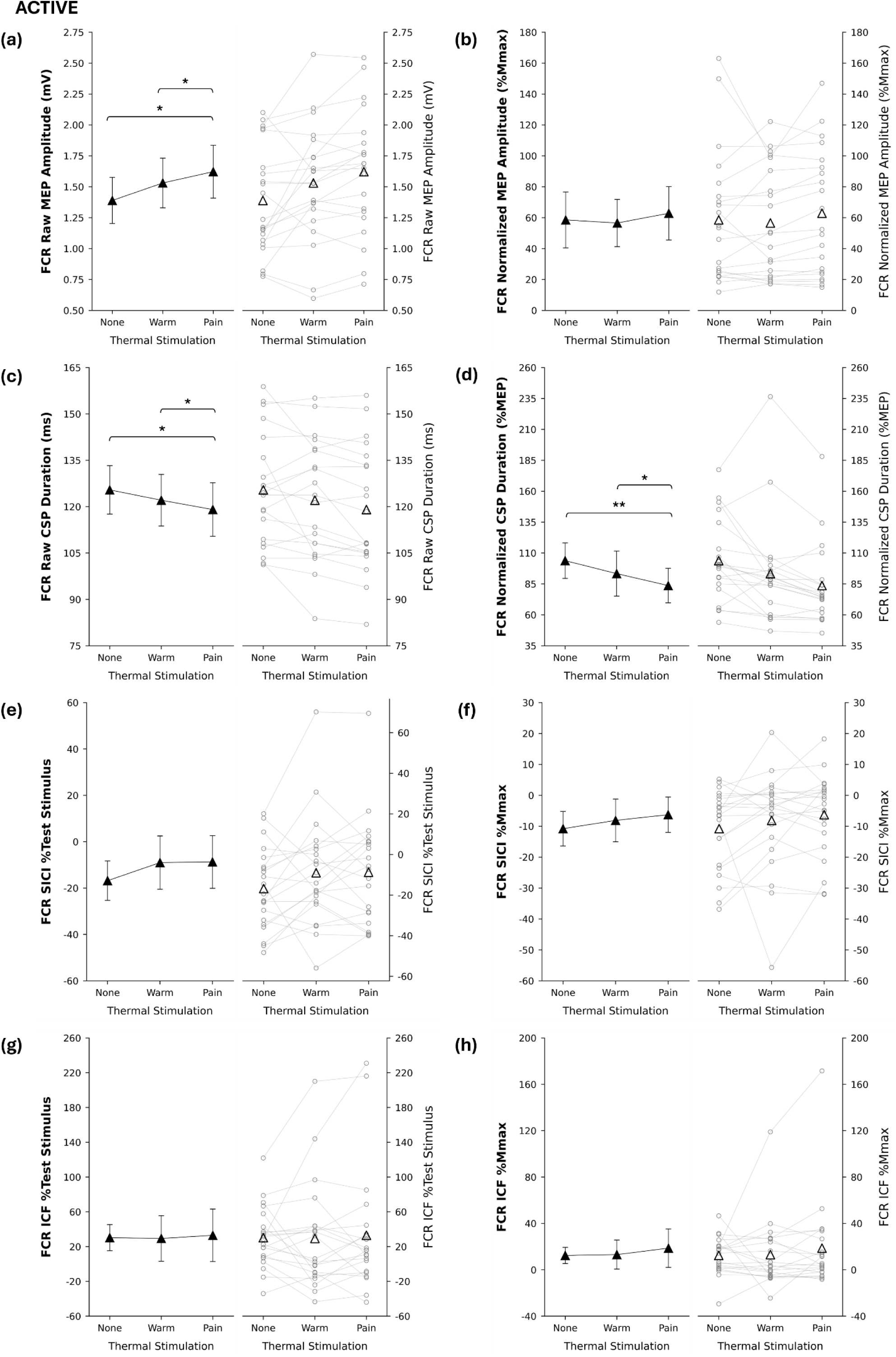
Effect of thermal stimulation on motor evoked potential (MEP), cortical silent period (CSP), short-interval intracortical inhibition (SICI), intracortical facilitation (ICF) recorded from the flexor carpi radialis (FCR) during voluntary isometric wrist flexions at 20% maximal voluntary contraction peak torque. In all panels, the left side shows mean values (black triangles) with 95% confidence interval error bars, while the right side shows individual data points (gray circles) along with the mean (black triangles). *Panel (a)* shows raw MEP amplitude expressed in millivolt (mV). *Panel (b)* shows MEP amplitudes normalized by Mmax amplitude. *Panel (c)* shows raw CSP duration expressed in milliseconds (ms). *Panel (d)* shows CSP duration normalised by Mmax amplitude. *Panel (e)* shows SICI expressed in percentage of the test stimulus. *Panel (f)* shows SICI expressed in percentage of the Mmax. *Panel (g)* shows ICF expressed in percentage of the test stimulus. *Panel (h)* shows ICF expressed in percentage of Mmax. * and ** for difference between two temperatures with p <.05 and p <.01, respectively.

#### CSP

During voluntary wrist flexions, mean raw CSP duration was 125.42 ± 18.31 ms for control condition, 122.08 ± 19.51 ms for warm condition and 119.05 ± 20.23 ms for pain condition. Mean normalized CSP duration was 101.31 ± 34.75 for control condition, 92.16 ± 41.77 for warm condition and 82.73 ± 32.12 for pain condition. There was a main effect of temperature on raw CSP duration (*F*(1.40, 26.65) = 5.58, *p* =.017, *η²_p_* =.227 [.007,.444]; **Fig. 5c**). Post hoc comparisons revealed that CSP duration was reduced during pain compared to control (*t*(19) = - 3.03, *p* =.020, d =-0.329 [-0.646,-0.012]) and pain compared to warm conditions (*t*(19) =-2.65, *p* =.032, *d* =-0.156 [-0.325, 0.012]), with no difference between control and warm conditions (*t*(19) = 1.46, *p* =.159, *d* = 0.173 [-0.145, 0.490]). There was a main effect of temperature on CSP duration normalized to MEP amplitude (*F*(1.43, 28.71) = 7.00, *p* =.007, *η²_p_* =.259 [.024,.466]; **Fig. 5d**). Post hoc comparisons revealed that CSP/MEP was reduced during pain compared to control (*t*(20) =-3.56, *p* =.006, *d* =-0.510 [-0.940,-0.080]) and pain compared to warm conditions (*t*(20) =-2.96, *p* =.015, *d* =-0.259 [-0.511, 0.007]), but not between control and warm conditions (*t*(20) = 1.51, *p* =.146, *d* = 0.251 [-0.194, 0.697]).

#### TS

During voluntary wrist flexions, mean test stimulus amplitude was 1.21 ± 0.32 mV for control condition, 1.24 ± 0.30 mV for warm condition and 1.23 ± 0.34 mV for pain condition. No main effect of temperature was found on active FCR test stimulus amplitude (*F*(1.95, 39.05) = 0.48, *p* =.619, *η²_p_* =.023 [.000,.140]).

#### SICI

During voluntary wrist flexions, mean SICI as percentage of test stimulus was-16.81 ± 19.95 % for control condition,-9.01 ± 26.86 % for warm condition and-8.74 ± 26.55 % for pain condition. Mean SICI as percentage of Mmax was-10.79 ± 13.07 % for control condition,-8.12 ± 16.18 % for warm condition and-6.26 ± 13.36 % for pain condition. No main effect of temperature was observed on SICI expressed as a percentage of the test stimulus (*F*(1.60, 31.94) = 2.00, *p* =.159, *η²_p_* =.091 [.000,.280]; **Fig. 5e**), or as a percentage of Mmax (*F*(1.70, 34.08) = 1.33, *p* =.274, *η²_p_* =.062 [.000,.231]; **Fig. 5f**).

#### ICF

During voluntary wrist flexions, mean ICF as percentage of test stimulus was 30.26 ± 35.18 % for control condition, 29.37 ± 62.32 % for warm condition and 32.98 ± 70.39 % for pain condition. Mean ICF as percentage of Mmax was 12.36 ± 16.29 % for control condition, 13.07 ± 29.25 % for warm condition and 18.67 ± 38.81 % for pain condition. No main effect of temperature was observed on ICF expressed as percentage of the test stimulus (*F*(1.43, 28.62) = 0.07, *p* =.881, *η²_p_* =.003 [.000,.097]; **Fig. 5g**) or as percentage of Mmax (*F*(1.26, 24.02) = 0.74, *p* =.429, *η²_p_* =.037 [.000,.236]; **Fig. 5h**).

#### Extensor carpi radialis

While the hotspot and stimulation intensity were determined specifically for the FCR muscle, we also collected for exploratory analyses responses in the ECR muscle. Detailed results are presented in supplementary material 3. Briefly, during voluntary wrist flexion, we did not observe any effect of temperature on MEPs amplitude, CSP duration, SICI and ICF (all *ps* >.175, all *η²_p_* <.089).

### Repeated measures correlation

After adjusting for multiple comparisons (*n* = 6), repeated-measures correlation analyses revealed a moderate positive correlation between temperature perception and effort perception (*r_rm_*(41) =.48, 95% CI [.21,.68], *p_adj_* =.006), indicating that higher perception of the stimulation temperature was associated with increased perception of effort. The remaining variables did not reach statistical significance with or without Holm–Bonferroni correction. Detailed results for all repeated-measures correlations are provided in **Table 1**.

**Table 1:** Repeated-measures correlations between perception of effort ratings and subjective or neurophysiological variables. . Cortical silent period (CSP), flexor carpi radialis (FCR), intracortical facilitation (ICF), motor evoked potential (MEP), short-interval intracortical inhibition (SICI), test stimulus (TS). ***_!_*** Note: Adjusted p-values marked with an asterisk were set equal to the previous rank to satisfy the principle of monotonicity.

| *Variable* | *df* | *r<sub>rm</sub> [95%CI]* | *Uncorrected p* | *Holm-Bonferroni p<sub>adj</sub>* |
| --- | --- | --- | --- | --- |
| **Temperature Perception** | 41 | .48 [.21, .68] | .001 | **<u>.006</u>** |
| Temperature Intensity | 41 | .29 [-.01, .54] | .06 | .300 |
| FCR CSP normalized | 39 | -.23 [-.51, .08] | .14 | .560 |
| FCR SICI %TS | 41 | -.19 [-.46, .12] | .233 | .699 |
| FCR MEP normalized | 41 | .18 [-.13, .45] | .252 | .699<sup>!</sup> |
| FCR ICF %TS | 40 | .17 [-.15, .45] | .293 | .699<sup>!</sup> |

## Discussion

The present study investigated whether contralateral pain reduces corticospinal and intracortical excitability during submaximal isometric wrist flexions, thereby contributing to the increased perceived effort associated with pain. Transcranial magnetic stimulation-derived neurophysiological measures were recorded from the right FCR and ECR muscles, while pain was induced on the left forearm. As the study focused a priori on the FCR muscle, TMS parameters were optimized for this muscle, and analyses involving the ECR muscle were considered exploratory.

Contralateral thermal stimulation effectively induced pain and increased perceived effort compared with the control and non-painful conditions, confirming that pain imposes an additional cost on task execution. Contrary to our initial hypotheses, pain did not reduce corticospinal or intracortical excitability. Instead, contralateral pain increased raw MEP amplitude and reduced CSP duration in the FCR muscle during voluntary contraction. Exploratory analysis of the ECR revealed that contralateral pain increased MEP amplitude and reduced SICI at rest. Repeated-measures correlation analyses further showed that higher pain perception, but not stimulus temperature intensity, was associated with greater effort perception.

### Pain effect on corticospinal and intracortical excitability

Two recent meta-analyses reported an inhibitory effect of experimental pain on corticospinal excitability at rest (Chowdhury et al., 2022; Rohel et al., 2021). However, most of the included studies applied pain to the hand and recorded MEPs from muscles directly exposed to the nociceptive stimulus, limiting the generalizability of findings to other anatomical sites. When experimental pain, regardless of the method used to induce pain, is applied contralateral to the recorded muscle at rest, studies have reported either no change in MEP amplitudes recorded during pain application (Alhassani et al., 2019; Le Pera et al., 2001; Winther et al., 2025) or increased MEP amplitude during (Delahunty et al., 2019) and 600ms after pain application (Algoet et al., 2018), providing no evidence for an inhibitory effect. Our results are in line with this literature, as we observed no inhibitory effect of contralateral pain on corticospinal excitability at rest.

Given the limited evidence regarding the effects of experimental pain on intracortical excitability at rest, studies using both remote and local pain models should be considered. Contralateral cold pain reduced SICI and increased ICF recorded from ECR muscle (Delahunty et al., 2019). Local chemical cutaneous hand pain decreased SICI without affecting ICF in the abductor pollicis brevis muscle (Fierro et al., 2010), whereas local chemical muscle hand pain did not alter SICI but reduced ICF in the first dorsal interosseous muscle (Schabrun & Hodges, 2012). In the present study, contralateral thermal heat pain did not alter intracortical excitability in the FCR muscle. These discrepancies across studies may be explained by differences in pain protocols and pain stimulus location.

During voluntary contraction, chemically induced cutaneous or muscle pain applied to the active muscle has been associated with reduced corticospinal excitability (Cheong et al., 2003; Martin et al., 2007). To our knowledge, the present study is the first to examine the effects of contralateral heat pain on corticospinal excitability during voluntary contraction. During isometric wrist flexions at 20% of maximal torque, our a priori analysis suggested increased FCR corticospinal excitability under contralateral pain.

Under painful stimulation, force output, raw MEP amplitude and background EMG increased, whereas CSP duration decreased. The disappearance of MEP and EMG differences after normalization to Mmax suggests that the increased raw MEP amplitude may be attributable to altered peripheral excitability. In lower limbs, nociceptive afferents can modulate motoneurons controlling the contralateral limb through a phenomenon called crossed-extension reflex (Sherrington, 1910), although descending cortical inputs can influence these spinal interactions (Rustamov et al., 2019). In the upper limbs, evidence for such crossed spinal modulation remains limited. Studies measuring upper limb H-reflex or M-wave during painful stimulation reported no changes in these measures (Le Pera et al., 2001; Svensson et al., 2003). Given the limited evidence on the influence of contralateral heat pain on peripheral measures, we cannot exclude the possibility that the observed increase in raw MEP amplitudes may reflect central mechanisms, particularly given the concurrent reduction in CSP duration. The reduction in CSP duration likely reflect a combination of two factors: (i) a slightly elevated voluntary drive (Matsugi, 2019), and (ii) reduced GABA_B_-mediated intracortical inhibition associated with pain (Butler et al., 2012).

To our knowledge, this is the first study to examine intracortical excitability during voluntary muscle contraction during contralateral heat pain. We did not observe any modulation of intracortical excitability. During active contraction, the motoneuron pool operates closer to firing threshold, requiring fewer cortical volleys to generate responses, reducing the conditioning stimuli capacity to modulate intracortical interneurons (Neige et al., 2020; Zoghi et al., 2003). Under such conditions, intracortical measures are likely less sensitive to detecting pain-related effects during voluntary muscle activity.

### Exploratory analysis on ECR

At rest, contralateral heat pain increased both raw and normalized MEP amplitudes and reduced SICI expressed as test stimulus percentage, suggesting reduced intracortical inhibition. Previous studies have shown that contralateral pain can reduce interhemispheric inhibition (Alhassani et al., 2019; Schabrun et al., 2016), a mechanism that normally contributes to lateralized motor control and limits mirror activity (Carson, 2020). In our study, pain applied to the left forearm may have reduced interhemispheric inhibition exerted by the right hemisphere, which received nociceptive input, onto the left motor cortex targeted by TMS. This putative reduction in interhemispheric inhibition may reflect a release of the non-pain-affected cortex from inhibitory control, which could facilitate protective motor strategies in the non-painful limb (Wiech & Tracey, 2013). However, ECR results should be interpreted with caution, as the TMS hotspot location and stimulation intensity were optimized for the FCR, our a priori muscle of interest.

### Is effort perception during pain related to corticospinal and intracortical excitability?

To examine whether effort perception was associated with pain-related modification in corticospinal and intracortical excitability, we conducted repeated-measures correlations between ratings of perceived effort and TMS-derived neurophysiological variables. Previous meta-analyses have reported inhibitory effects of experimental pain on these variables (Chowdhury et al., 2022; Rohel et al., 2021), suggesting that greater perceived effort during painful tasks might reflect the need to increase central motor command to overcome pain-related inhibition. However, the present findings, obtained in the context of contralateral heat pain, do not support a generalized inhibitory effect of pain, either at rest or during voluntary contraction. We did not observe any correlation between neurophysiological variables and ratings of perceived effort. In contrast, perceived effort was positively correlated with the perceived temperature but not with the actual temperature of the thermal stimulation. This results is consistent with the findings of the study of Mangin et al. (2026) in which computational analyses showed that perceived effort during cognitive and visuomotor task execution under contralateral thermal pain was better predicted by pain perception than by the actual temperature of stimulation. Taken together, these results suggest that the subjective experience of pain contributes more strongly to the increased effort perception experience than the nociceptive input alone. Furthermore, previous work has shown that, during a self-regulated motor output task in the presence of contralateral heat pain, greater force is produced for the same level of perceived effort, suggesting that perceived effort is not determined exclusively by the central motor command (O’Malley et al., 2026). By integrating these results with those of Monti et al. (2026), who reported altered SMA and MCC activity during a similar visuomotor task executed under pain, we propose that maintaining task performance reflects altered central resource allocation, such as increased cognitive control to sustain task engagement despite contralateral heat pain.

### Limitation and future perspective

Pain type and its anatomical location may differentially influence corticospinal and intracortical excitability. Our findings are specific to contralateral heat pain and to measurements of corticospinal and intracortical excitability in the forearm muscles, and cannot be generalized to other pain modalities or muscle groups. We encourage future studies to assess different muscles using the same experimental pain protocol, as well as the same muscle using different pain protocols or anatomical locations. In the present study, TMS parameters were optimized only for the FCR muscle. As we observed different responses between the FCR and ECR muscles, future studies should a priori focus on the ECR to better disentangle the differential effect of contralateral heat pain on the motor control sustaining wrist flexions. Additionally, Mmax was used as a peripheral measure of muscle excitability, and our study did not include measures of spinal excitability. As nociceptive fibers project not only at cortical level but also at spinal level, future research should incorporate indices of spinal excitability, such as cervicomedullary MEPs or H-reflex, to directly assess the influence of contralateral pain at the spinal level.

## Conclusion

Pain captures attention (Torta et al., 2017) and triggers pain-avoidance responses (Wiech & Tracey, 2013), increasing task difficulty. Consequently, behavioral regulation and cognitive control are required to perform tasks under pain (Monti et al., 2026; Seminowicz, 2007). As increased perceived effort during contralateral heat pain was not associated with inhibition of corticospinal or intracortical excitability, but was related only to the subjective experience of pain, the present findings support this view.

## Acknowledgements

We would like to thank Dr. Catherine Mercier for her feedback on the manuscript.

## Data availability

The dataset generated and analyzed during the current study is available at https://osf.io/fzxw3.

## Use of Generative AI Tools

The authors used ChatGPT (OpenAI) to assist with English language editing and grammar improvement during manuscript preparation. After using this tool, the authors reviewed and edited the content as needed and takes full responsibility for the content of the published article. All scientific content, data interpretation, and conclusions were generated solely by the authors.

## Author contributions

Author contributions are reported according to the CRediT author statement.

**Ilaria Monti:** Conceptualization, Methodology, Software, Validation, Formal analysis, Investigation, Data Curation, Writing - Original Draft, Visualization, Project administration. **Maxime Bergevin:** Methodology, Software, Validation, Writing - Review & Editing. **Madhumitha Murugavel Sangeetha:** Investigation, Data Curation, Writing - Review & Editing. **Thomas Mangin:** Conceptualization, Methodology, Software, Writing - Review & Editing. **Jason Neva:** Conceptualization, Methodology, Resources, Writing - Review & Editing. **Mathieu Roy:** Resources, Writing - Review & Editing. **Pierre Rainville:** Conceptualization, Methodology, Resources, Writing - Review & Editing, Supervision. **Benjamin Pageaux:** Conceptualization, Methodology, Validation, Resources, Visualization, Writing - Review & Editing, Supervision, Funding acquisition.

## References

Algoet, M., Duque, J., Iannetti, G. D., & Mouraux, A. (2018). Temporal Profile and Limb-specificity of Phasic Pain-Evoked Changes in Motor Excitability. Neuroscience, 386, 240–255. 10.1016/j.neuroscience.2018.06.039

Alhassani, G., Liston, M. B., & Schabrun, S. M. (2019). Interhemispheric Inhibition Is Reduced in Response to Acute Muscle Pain: A Cross-Sectional Study Using Transcranial Magnetic Stimulation. Journal of Pain, 20(9), 1091–1099. 10.1016/j.jpain.2019.03.007

Amador, N., & Fried, I. (2004). Single-neuron activity in the human supplementary motor area underlying preparation for action. Journal of Neurosurgery, 100(2), 250–259. 10.3171/JNS.2004.100.2.0250

Badawy, R. A. B., Loetscher, T., Macdonell, R. A. L., & Brodtmann, A. (2012). Cortical excitability and neurology: Insights into the pathophysiology. Functional Neurology, 27(3).

Bergevin, M., Steele, J., Payen de la Garanderie, M., Feral-Basin, C., Marcora, S. M., Rainville, P., Caron, J. G., & Pageaux, B. (2023). Pharmacological Blockade of Muscle Afferents and Perception of Effort: A Systematic Review with Meta-analysis. Sports Medicine, 53(2), 415–435. 10.1007/S40279-022-01762-4

Botvinick, M. M., Carter, C. S., Braver, T. S., Barch, D. M., & Cohen, J. D. (2001). Conflict monitoring and cognitive control. Psychological Review, 108(3), 624–652. 10.1037/0033-295X.108.3.624

Butler, J. E., Petersen, N. C., Herbert, R. D., Gandevia, S. C., & Taylor, J. L. (2012). Origin of the low-level EMG during the silent period following transcranial magnetic stimulation. Clinical Neurophysiology, 123(7), 1409–1414. 10.1016/j.clinph.2011.11.034

Byström, S. E. G., & Kilbom, Å. (1990). Physiological response in the forearm during and after isometric intermittent handgrip. European Journal of Applied Physiology and Occupational Physiology, 60(6), 457–466. 10.1007/BF00705037/METRICS

Cancela, T., Gendolla, G. H. E., & Silvestrini, N. (2023). Pain and gain: Monetary incentive moderates pain’s impact on effort-related cardiac response. Psychophysiology, 60. 10.1111/psyp.14231

Carson, R. G. (2020). Inter-hemispheric inhibition sculpts the output of neural circuits by co-opting the two cerebral hemispheres. Journal of Physiology, 598(21), 4781–4802. 10.1113/JP279793

Cheong, J. Y., Yoon, T. S., & Lee, S. J. (2003). Evaluations of inhibitory effect on the motor cortex by cutaneous pain via application of capsaicin. Electromyography and Clinical Neurophysiology, 43(4), 203–210. https://europepmc.org/article/med/12836584

Chowdhury, N. S., Chang, W. J., Millard, S. K., Skippen, P., Bilska, K., Seminowicz, D. A., & Schabrun, S. M. (2022). The Effect of Acute and Sustained Pain on Corticomotor Excitability: A Systematic Review and Meta-Analysis of Group and Individual Level Data. The Journal of Pain, 23(10), 1680–1696. 10.1016/J.JPAIN.2022.04.012

De Morree, H. M., & Marcora, S. M. (2015). Psychobiology of perceived effort during physical tasks. In Handbook of Biobehavioral Approaches to Self-Regulation (pp. 255–270). Springer New York. 10.1007/978-1-4939-1236-0_17

Delahunty, E. T., Bisset, L. M., & Kavanagh, J. J. (2019). Intracortical motor networks are affected in both the contralateral and ipsilateral hemisphere during single limb cold water immersion. Experimental Physiology, 104(8), 1296–1305. 10.1113/EP087745

Dum, R. P., Levinthal, D. J., & Strick, P. L. (2009). The spinothalamic system targets motor and sensory areas in the cerebral cortex of monkeys. Journal of Neuroscience, 29(45), 14223–14235. 10.1523/JNEUROSCI.3398-09.2009

El Bouse, A. O., Gabriel, D. A., & Tokuno, C. D. (2013). Examining the reliability of the flexor carpi radialis V-wave at different levels of muscle contraction. Journal of Electromyography and Kinesiology, 23(2), 296–301. 10.1016/J.JELEKIN.2012.10.008

Fierro, B., De Tommaso, M., Giglia, F., Giglia, G., Palermo, A., & Brighina, F. (2010). Repetitive transcranial magnetic stimulation (rTMS) of the dorsolateral prefrontal cortex (DLPFC) during capsaicin-induced pain: Modulatory effects on motor cortex excitability. Experimental Brain Research, 203(*1*), 31–38. 10.1007/s00221-010-2206-6

Forman, D. A., Forman, G. N., Mugnosso, M., Zenzeri, J., Murphy, B., & Holmes, M. W. R. (2020). Sustained Isometric Wrist Flexion and Extension Maximal Voluntary Contractions Similarly Impair Hand-Tracking Accuracy in Young Adults Using a Wrist Robot. Frontiers in Sports and Active Living, 2, 53. 10.3389/fspor.2020.00053

Gillies, M. J., Huang, Y., Hyam, J. A., Aziz, T. Z., & Green, A. L. (2019). Direct neurophysiological evidence for a role of the human anterior cingulate cortex in central command. Autonomic Neuroscience: Basic and Clinical, 216. 10.1016/j.autneu.2018.09.004

Gueugneau, N., Martin, A., Gaveau, J., & Papaxanthis, C. (2023). Gravity-efficient motor control is associated with contraction-dependent intracortical inhibition. IScience, 26(7), 107150. 10.1016/j.isci.2023.107150

Kujirai, T., Caramia, M. D., Rothwell, J. C., Day, B. L., Thompson, P. D., Ferbert, A., Wroe, S., Asselman, P., & Marsden, C. D. (1993). Corticocortical inhibition in human motor cortex. The Journal of Physiology, 471(1), 501–519. 10.1113/JPHYSIOL.1993.SP019912

Lackmy, A., & Marchand-Pauvert, V. (2010). The estimation of short intra-cortical inhibition depends on the proportion of spinal motoneurones activated by corticospinal inputs. Clinical Neurophysiology, 121(4), 612–621. 10.1016/j.clinph.2009.12.011

Le Pera, D., Graven-Nielsen, T., Valeriani, M., Oliviero, A., Di Lazzaro, V., Tonali, P. A., & Arendt-Nielsen, L. (2001). Inhibition of motor system excitability at cortical and spinal level by tonic muscle pain. Clinical Neurophysiology, 112(9), 1633–1641. 10.1016/S1388-2457(01)00631-9

Legrain, V., Damme, S. Van, Eccleston, C., Davis, K. D., Seminowicz, D. A., & Crombez, G. (2009). A neurocognitive model of attention to pain: behavioral and neuroimaging evidence. Pain, 144(3), 230–232. 10.1016/J.PAIN.2009.03.020

Mangin, T., Monti, I., Marcotte, M., Baudry, S., Roy, M., Rainville, P., & Pageaux, B. (2026). Maintaining performance under pain is effortful: experimental and computational evidence. BioRxiv, 2026.02.13.705857. 10.64898/2026.02.13.705857

Mangin, T., & Pageaux, B. (2026). Effort and its perception revisited: How physical-domain insights could lead toward a unified theory. *Cognitive, Affective*, & Behavioral Neuroscience, 1–47. 10.3758/s13415-026-01411-7

Martin, P. G., Weerakkody, N., Gandevia, S. C., & Taylor, J. L. (2007). Group III and IV muscle afferents differentially affect the motor cortex and motoneurones in humans. The Journal of Physiology, 586(Pt 5), 1277. 10.1113/JPHYSIOL.2007.140426

Matsugi, A. (2019). Changes in the cortical silent period during force control. Somatosensory & Motor Research, 36(1), 8–13. 10.1080/08990220.2018.1563536

Monti, I., Picard, M.-E., Mangin, T., Bergevin, M., Gruet, M., Baudry, S., Otto, A. R., Chen, J.-I., Roy, M., Rainville, P., & Pageaux, B. (2026). Facing pain is effortful: key role of the supplementary motor area and anterior midcingulate cortex. BioRxiv, 2026.04.17.719211. 10.64898/2026.04.17.719211

Neige, C., Grosprêtre, S., Martin, A., & Lebon, F. (2020). Influence of voluntary contraction level, test stimulus intensity and normalization procedures on the evaluation of short-interval intracortical inhibition. Brain Sciences, 10(7), 1–16. 10.3390/BRAINSCI10070433

Ni, Z., & Chen, R. (2012). Intracortical circuits and their interactions in human primary motor cortex. In Cortical Connectivity: Brain Stimulation for Assessing and Modulating Cortical Connectivity and Function. 10.1007/978-3-642-32767-4_3

O’Malley, C., Mangin, T., Bergevin, M., Monti, I., Fullerton, C., Mauger, A., Rainville, P., & Pageaux, B. (2026). Experimental thermal pain and naturally occurring muscle pain have different effects on force production during a fixed perceived effort handgrip task. 10.31234/osf.io/fk48y_v2

Opie, G. M., & Semmler, J. G. (2014). Modulation of short-and long-interval intracortical inhibition with increasing motor evoked potential amplitude in a human hand muscle. Clinical Neurophysiology, 125(7), 1440–1450. 10.1016/j.clinph.2013.11.015

Ortu, E., Deriu, F., Suppa, A., Tolu, E., & Rothwell, J. C. (2008). Effects of volitional contraction on intracortical inhibition and facilitation in the human motor cortex. The Journal of Physiology, 586(21), 5147–5159. 10.1113/JPHYSIOL.2008.158956

Pageaux, B., Angius, L., Hopker, J. G., Lepers, R., & Marcora, S. M. (2015). Central alterations of neuromuscular function and feedback from group III-IV muscle afferents following exhaustive high-intensity one-leg dynamic exercise. American Journal of Physiology - Regulatory Integrative and Comparative Physiology, 308(12)(PG-R1008-R1020), R1008–R1020. 10.1152/ajpregu.00280.2014

Preston, J., & Wegner, D. M. (2009). Elbow grease: when action feels like work BT - Oxford Handbook of Human Action. In Oxford Handbook of Human Action (Issue 27, pp. 469–486).

Richter, M., Gendolla, G. H. E., & Wright, R. A. (2016). Three Decades of Research on Motivational Intensity Theory: What We Have Learned About Effort and What We Still Don’t Know. Advances in Motivation Science, 3, 149–186. 10.1016/BS.ADMS.2016.02.001

Rohel, A., Bouffard, J., Patricio, P., Mavromatis, N., Billot, M., Roy, J. S., Bouyer, L., Mercier, C., & Masse-Alarie, H. (2021). The effect of experimental pain on the excitability of the corticospinal tract in humans: A systematic review and meta-analysis. European Journal of Pain, 25(6), 1209–1226. 10.1002/EJP.1746

Rossini, P. M., Barker, A. T., Berardelli, A., Caramia, M. D., Caruso, G., Cracco, R. Q., Dimitrijević, M. R., Hallett, M., Katayama, Y., Lücking, C. H., Maertens de Noordhout, A. L., Marsden, C. D., Murray, N. M. F., Rothwell, J. C., Swash, M., & Tomberg, C. (1994). Non-invasive electrical and magnetic stimulation of the brain, spinal cord and roots: basic principles and procedures for routine clinical application. Report of an IFCN committee. Electroencephalography and Clinical Neurophysiology, 91(2), 79–92. 10.1016/0013-4694(94)90029-9

Rossini, P. M., Burke, D., Chen, R., Cohen, L. G., Daskalakis, Z., Di Iorio, R., Di Lazzaro, V., Ferreri, F., Fitzgerald, P. B., George, M. S., Hallett, M., Lefaucheur, J. P., Langguth, B., Matsumoto, H., Miniussi, C., Nitsche, M. A., Pascual-Leone, A., Paulus, W., Rossi, S.,…Ziemann, U. (2015). Non-invasive electrical and magnetic stimulation of the brain, spinal cord, roots and peripheral nerves: Basic principles and procedures for routine clinical and research application. An updated report from an I.F.C.N. Committee. Clinical Neurophysiology, 126(6), 1071–1107. 10.1016/J.CLINPH.2015.02.001

Rustamov, N., Northon, S., Tessier, J., Leblond, H., & Piché, M. (2019). Integration of bilateral nociceptive inputs tunes spinal and cerebral responses. Scientific Reports 2019 9:1, 9(1), 7143-. 10.1038/s41598-019-43567-y

Schabrun, S. M., Christensen, S. W., Mrachacz-Kersting, N., & Graven-Nielsen, T. (2016). Motor Cortex Reorganization and Impaired Function in the Transition to Sustained Muscle Pain. Cerebral Cortex, 26(5), 1878–1890. 10.1093/cercor/bhu319

Schabrun, S. M., & Hodges, P. W. (2012). Muscle Pain Differentially Modulates Short Interval Intracortical Inhibition and Intracortical Facilitation in Primary Motor Cortex. The Journal of Pain, 13(2), 187–194. 10.1016/J.JPAIN.2011.10.013

Seminowicz, D. A. (2007). Functional MRI studies of pain and cognition interactions. Dissertation Abstracts International: Section B: The Sciences and Engineering, 68(6-B PG-3595), 3595. https://ovidsp.ovid.com/ovidweb.cgi?T=JS&CSC=Y&NEWS=N&PAGE=fulltext&D=psyc 5&AN=2007-99240-061 NS-

Shanks, M. J., Cirillo, J., Stinear, C. M., & Byblow, W. D. (2023). Reliability of a TMS-derived threshold matrix of corticomotor function. Experimental Brain Research 2023 241:11, 241(11), 2829–2843. 10.1007/S00221-023-06725-3

Sherrington, C. S. (1910). Flexion-reflex of the limb, crossed extension-reflex, and reflex stepping and standing. The Journal of Physiology, 40(1–2), 28. 10.1113/jphysiol.1910.sp001362

Stagg, C. J., Bestmann, S., Constantinescu, A. O., Moreno Moreno, L., Allman, C., Mekle, R., Woolrich, M., Near, J., Johansen-Berg, H., & Rothwell, J. C. (2011). Relationship between physiological measures of excitability and levels of glutamate and GABA in the human motor cortex. The Journal of Physiology, 589(Pt 23), 5845. 10.1113/JPHYSIOL.2011.216978

Svensson, P., Miles, T. S., McKay, D., & Ridding, M. C. (2003). Suppression of motor evoked potentials in a hand muscle following prolonged painful stimulation. *European Journal of Pain (London*, England), 7(1), 55–62. 10.1016/S1090-3801(02)00050-2

Torta, D. M., Legrain, V., Mouraux, A., & Valentini, E. (2017). Attention to pain! A neurocognitive perspective on attentional modulation of pain in neuroimaging studies. Cortex; a Journal Devoted to the Study of the Nervous System and Behavior, 89, 120–134. 10.1016/J.CORTEX.2017.01.010

Wiech, K., & Tracey, I. (2013). Pain, decisions, and actions: a motivational perspective. Frontiers in Neuroscience, *7*(7 APR). 10.3389/FNINS.2013.00046

Winther, B. H., Arendt-Nielsen, L., Graugaard, J. B., Jespersen, H. G., Elberling, J., Martino, E. De, & Vecchio, S. Lo. (2025). Experimental acute muscle pain and itch induce similar inhibitory effects on corticospinal excitability. Pain Reports, 10(6), e1341. 10.1097/PR9.0000000000001341

Zoghi, M., Pearce, S. L., & Nordstrom, M. A. (2003). Differential modulation of intracortical inhibition in human motor cortex during selective activation of an intrinsic hand muscle. The Journal of Physiology, 550(Pt 3), 933–946. 10.1113/JPHYSIOL.2003.042606

