## Supplementary materials for "Increased perceived effort during contralateral thermal heat pain is not explained by increased intracortical and corticospinal inhibition"

Because the peripheral nerve stimulation site and intensity were optimized for the flexor carpi radialis (FCR), the M-wave values reported for the extensor carpi radialis (ECR) are likely submaximal and should not be interpreted as true maximal M-waves, unlike those obtained for the FCR, our a priori muscle of interest. However, as part of the exploratory ECR analyses, we report M-wave amplitudes and provide M-wave-normalized values to account, as far as possible, for potential peripheral changes across conditions. Readers should therefore interpret ECR variables normalized to the M-wave with caution.

### Supplementary material 1

#### Extensor carpi radialis Mwave during rest and active session

For ECR at rest, mean Mwave amplitude was 1.09 ± 0.96 mV for control condition, 1.11 ± 0.99 mV for warm condition and 1.11 ± 1.0 mV for pain condition. ECR Mwave amplitude at rest did not change with temperature ($F(1.83, 33.07) = 0.207$, $p = .796$, $\eta^2_p = .01$ [.000, .115]).

During active session, for ECR the mean Mwave amplitude was 1.12 ± 0.76 mV for control condition, 1.07 ± 0.8 mV for warm condition and 1.08 ± 0.76 mV for pain condition. No main effect of temperature was found on Mwave amplitude during voluntary wrist flexions ($F(1.63, 32.68) = 2.295$, $p = .125$, $\eta^2_p = .103$ [.000, .292]) muscles.

#### Background EMG rest session

*FCR MEP.* During rest, the temperature of stimulation did not alter the raw RMS of the background EMG ($F(1.09, 20.62) = 0.007$, $p = .948$, $\eta^2_p = .001$ [.000, .063]; **Table S1**), even when normalized to Mwave ($F(1.10, 20.90) = 0.038$, $p = .869$, $\eta^2_p = .002$ [.000, .128]; **Table S1**).

*FCR SICI.* During rest, no main effect of temperatures was found on the raw RMS of the background EMG ($F(1.13, 21.38) = 1.964$, $p = .175$, $\eta^2_p = .094$ [.000, .337]; **Table S1**). When normalized to the Mwave, an effect of temperature was found on the RMS of the background EMG ($F(1.07, 20.23) = 4.268$, $p = .050$, $\eta^2_p = .183$ [.000, .438]; **Table S1**). However, post hoc comparisons revealed no significant differences between conditions (control vs. warm: $t(19) = 2.15$, $p = .134$), $d = 0.341$ [-0.099, 0.782]; control vs. pain: $t(19) = 2.02$, $p = .134$, $d = 0.315$ [-0.116, 0.746]; warm vs. pain: $t(19) = -0.813$, $p = .427$, $d = -0.026$ [-0.113, 0.060]).

*FCR ICF.* During rest, no main effect of temperatures was found on the raw RMS of the background EMG ($F(1.17, 22.14) = 0.497$, $p = .516$, $\eta^2_p = .026$ [.000, .225]; **Table S1**), nor when normalized to the Mwave ($F(1.08, 20.48) = 3.028$, $p = .095$, $\eta^2_p = .137$ [.000, .391]; **Table S1**).

*ECR MEP.* During rest, the temperature of stimulation did not alter the raw RMS of the background EMG ($F(1.79, 26.79) = 1.043$, $p = .359$, $\eta^2_p = .065$ [.000, .252]; **Table S1**) even when normalized to Mwave ($F(1.62, 24.22) = 2.921$, $p = .083$, $\eta^2_p = .163$ [.000, .386]; **Table S1**).

*ECR SICI.* During rest, no main effect of temperatures was found on the raw RMS of the background EMG ($F$(1.13, 20.35) = 1.570, $p$ = .227, $\eta^2_p$ = .080 [.000, .325]; **Table S1**), nor on the normalized RMS ($F$(1.26, 22.70) = 2.602, $p$ = .114, $\eta^2_p$ = .126 [.000, .363]; **Table S1**).

*ECR ICF.* During rest, no main effect of temperatures was found on the raw RMS of the background EMG ($F$(1.26, 22.58) = 0.589, $p$ = .487, $\eta^2_p$ = .032 [.000, .232]; **Table S1**), nor when normalized to the Mwave ($F$(1.61, 28.95) = 0.738, $p$ = .459, $\eta^2_p$ = .039 [.000, .209]; **Table S1**).

| ***Muscle*** | ***TMS condition*** | ***Muscle state*** | ***RMS (mV) control*** | ***RMS (mV) warm*** | ***RMS (mV) pain*** | ***RMS (%Mwave) control*** | ***RMS (%Mwave) warm*** | ***RMS (%Mwave) pain*** |
|---|---|---|---|---|---|---|---|---|
| **FCR** | MEP | rest | 0.003 | 0.003 | 0.003 | 0.129 | 0.132 | 0.130 |
| **FCR** | SICI | rest | 0.004 | 0.003 | 0.003 | 0.174 | 0.128 | 0.132 |
| **FCR** | ICF | rest | 0.004 | 0.003 | 0.003 | 0.168 | 0.129 | 0.127 |
| **ECR** | MEP | rest | 0.007 | 0.007 | 0.007 | 1.055 | 1.199 | 1.269 |
| **ECR** | SICI | rest | 0.007 | 0.008 | 0.009 | 1.012 | 1.117 | 1.218 |
| **ECR** | ICF | rest | 0.008 | 0.009 | 0.008 | 1.182 | 1.166 | 1.336 |

**Table S 1: Mean values of RMS of the background EMG at rest expressed in mV and as percentage of Mwave amplitude.** No comparisons between temperatures reached significance. Extensor carpi radialis (ECR), flexor carpi radialis (FCR), intracortical facilitation (ICF), motor evoked potential (MEP), compound muscle action potentials (Mwave), millivolt (mV), root mean square (RMS), short-interval intracortical inhibition (SICI).

#### Background EMG active session

*FCR MEP.* During voluntary wrist flexions, a main effect of temperature was found on the raw RMS of the background EMG ($F$(1.42, 28.34) = 7.099, $p$ = .007, $\eta^2_p$ = .262 [.025, .470]). Post hoc comparisons revealed that RMS was higher during pain than in the control condition ($t$(20) = 3.46, $p$ = .007, $d$ = 0.215 [0.030, 0.401]; **Table S2**), with no difference between pain and warm conditions ($t$(20) = 0.82, $p$ = .422, $d$ = 0.033 [-0.073, 0.140]), and between warm and control conditions ($t$(20) = 2.38, $p$ = .054, $d$ = 0.182 [-0.031, 0.396]). When RMS was normalized to Mwave, the main effect of temperature disappeared ($F$(1.12, 22.34) = 0.483, $p$ = .515, $\eta^2_p$ = .024 [.000, .223]; **Table S2**).

*FCR SICI.* During voluntary wrist flexions, there was no main effect of temperature on the raw RMS of the background EMG ($F$(1.26, 25.10) = 0.185, $p$ = .727, $\eta^2_p$ = .009 [.000, .158]; **Table S2**), nor when normalized to the Mwave ($F$(1.26, 25.15) = 0.738, $p$ = .428, $\eta^2_p$ = .036 [.000, .229]; **Table S2**).

*FCR ICF.* During voluntary wrist flexions, there was no main effect of temperature for FCR muscle on the raw RMS of the background EMG ($F$(1.26, 25.19) = 0.162, $p$ = .748, $\eta^2_p$ = .008 [.000, .152]; **Table S2**), nor when normalized to Mwave ($F$(1.52, 28.90) = 0.044, $p$ = .919, $\eta^2_p$ = .002 [.000, .070]; **Table S2**).

*ECR MEP.* During voluntary wrist flexions, no main effect of temperature was observed for the ECR muscle on the raw RMS of the background EMG ($F$(1.13, 22.70) = 0.551, $p$ = .487, $\eta^2_p$ = .027 [.000, .227]; **Table S2**), nor when normalized by the Mwave ($F$(1.22, 24.41) = 0.210, $p$ = .699, $\eta^2_p$ = .010 [.000, .168]; **Table S2**).

*ECR SICI.* During voluntary wrist flexions, there was a main effect of temperature on the raw RMS EMG background ($F$(1.23, 24.56) = 8.756, $p$ = .004, $\eta^2_p$ = .304 [.037, .520]). Post hoc comparisons revealed that the RMS was lower during the warm compared to control condition ($t$(20) = -3.65, $p$ = .005, $d$ = -0.482 [-0.880, -0.084]; **Table S2**) and compared to pain condition ($t$(20) = -2.66, $p$ = .030, $d$ = -0.148 [-0.306, 0.010]). Post hoc comparisons revealed that the RMS was lower during the pain compared to control condition ($t$(20) = -2.29, $p$ = .033, $d$ = -0.334 [-0.738, 0.071]). When normalized to Mwave, the main effect of temperature disappeared ($F$(1.20, 24.02) = 2.926, $p$ = .094, $\eta^2_p$ = .128 [.000, .361]; **Table S2**).

*ECR ICF.* During voluntary wrist flexions, there was a main effect of temperature on the raw RMS EMG background ($F$(1.57, 31.47) = 12.04, $p$ = <.001, $\eta^2_p$ = .376 [.104, .554]). Post hoc comparisons revealed that the RMS was lower during the pain compared to control condition ($t$(20) = -3.531, $p$ = .004, $d$ = -0.480 [-0.887, -0.073]; **Table S2**) and during the warm compared to control condition ($t$(20) = -4.036, $p$ = .002, $d$ = -0.524 [-0.927, -0.122]; **Table S2**). No difference was found between the warm and pain conditions ($t$(20) = -0.537, $p$ = .597, $d$ = -0.044 [-0.260, 0.172]). When normalized to the Mwave, the main effect of temperature disappeared ($F$(1.36, 25.90) = 2.853, $p$ = .093, $\eta^2_p$ = .131 [.000, .352]; **Table S2**).

| ***Muscle*** | ***TMS condition*** | ***Muscle state*** | ***RMS (mV) control*** | ***RMS (mV) warm*** | ***RMS (mV) pain*** | ***RMS (%Mwave) control*** | ***RMS (%Mwave) warm*** | ***RMS (%Mwave) pain*** |
|---|---|---|---|---|---|---|---|---|
| **FCR** | MEP | active | **0.045** | 0.049 | **0.050** | 1.850 | 1.794 | 1.895 |
| **FCR** | SICI | active | 0.056 | 0.055 | 0.056 | 2.202 | 2.116 | 2.096 |
| **FCR** | ICF | active | 0.057 | 0.057 | 0.056 | 2.229 | 2.199 | 2.133 |
| **ECR** | MEP | active | 0.042 | 0.039 | 0.042 | 5.710 | 5.423 | 5.451 |
| **ECR** | SICI | active | **0.053** | **0.041** | **0.045** | 6.705 | 5.525 | 5.853 |
| **ECR** | ICF | active | **0.057** | **0.043** | **0.044** | 7.120 | 5.601 | 5.941 |

**Table S 2: Mean values of RMS of the background EMG during voluntary wrist flexions expressed in mV and as percentage of Mwave amplitude**. For MEP of FCR, the RMS was higher during pain compared to control condition. For SICI of ECR, the RMS was lower during warm compared to control condition. For ICF of ECR, the RMS was lower during pain and warm compared to control condition. No other comparisons reached significance. Extensor carpi radialis (ECR), flexor carpi radialis (FCR), intracortical facilitation (ICF), motor evoked potential (MEP), maximal compound muscle action potentials (Mwave), millivolt (mV), root mean square (RMS), short-interval intracortical inhibition (SICI).

*Force MEP*. There was a main effect of temperature on the mean force during the 100 ms preceding TMS pulses eliciting MEPs ($F$(1.80, 35.99) = 3.369, $p$ = .050, $\eta^2_p$ = .144 [.000, .330]). Post hoc comparisons revealed that participants produced 0.86% more force during pain (19.84 ± 0.26% MVC) compared to warm (19.67 ± 0.21% MVC) condition ($t$(20) = 3.13, $p$ = .016, $d$ = 0.679 [0.046, 1.311]), no significant difference was found between control (19.72 ± 0.27% MVC) and pain conditions ($t$(20)

= -1.56, $p$ = .320, $d$ = -0.444 [-1.207, 0.320]), and between control and warm conditions ($t(20)$ = 0.81, $p$ = .427, $d$ = 0.235 [-0.529, 0.999]).

*Force SICI*. No main effect of temperature was found on the mean force 100ms before the TMS pulse eliciting conditioned MEPs for SICI ($F(1.92, 38.22)$ = 0.311, $p$ = .725, $\eta^2_p$ = .015 [.000, .119]).

*Force ICF*. No main effect of temperature was found on the mean force 100ms before the TMS pulse eliciting conditioned MEPs for ICF ($F(1.99, 39.80)$ = 0.482, $p$ = .620, $\eta^2_p$ = .024 [.000, .401]).

### Supplementary material S2

#### Extensor carpi radialis (ECR) corticospinal and intracortical excitability at rest

*ECR MEP.* During rest, mean raw MEP amplitude was 0.58 ± 0.25 mV for control condition, 0.58 ± 0.26 mV for warm condition and 0.72 ± 0.33 mV for pain condition. Mean normalized MEP amplitude was 86.20 ± 57.14 for control condition, 92.46 ± 75.13 for warm condition and 121.37 ± 100.26 for pain condition. A significant main effect of temperature was found on the raw MEP amplitude ($F$(1.29, 19.35) = 6.623, $p$ = .013, $\eta^2_p$ = 0.306 [.016, .538]; **Fig. S1a**). Post hoc comparisons revealed an increased MEP amplitude during pain compared to warm ($t$(15) = 3.803, $p$ = .005, $d$ = 0.485 [0.067, 0.904]). No significant difference was found between control and warm conditions ($t$(15) = -0.080, $p$ = .937, $d$ = -0.009 [-0.330, 0.311]) and during control and pain conditions ($t$(15) = -2.412, $p$ = .058, $d$ = 0.495 [-1.098, 0.109]).). The observed effect remained significant when the MEP amplitude was normalized to the Mwave ($F$(1.16, 17.42) = 5.754, $p$ = .024, $\eta^2_p$ = .277 [.001, .527]; **Fig. S1b**). Post hoc comparisons revealed an increased MEP amplitude during pain compared to warm condition ($t$(15) = 3.516, $p$ = .009, $d$ = 0.364 [0.033, 0.695]). No significant difference was found between the control and warm conditions ($t$(15) = -0.732, $p$ = .476, $d$ = -0.079 [-0.371, 0.214]) and between control and pain conditions ($t$(15) = -2.338, $p$ = .067, $d$ = -0.442 [-0.997, 0.112]).

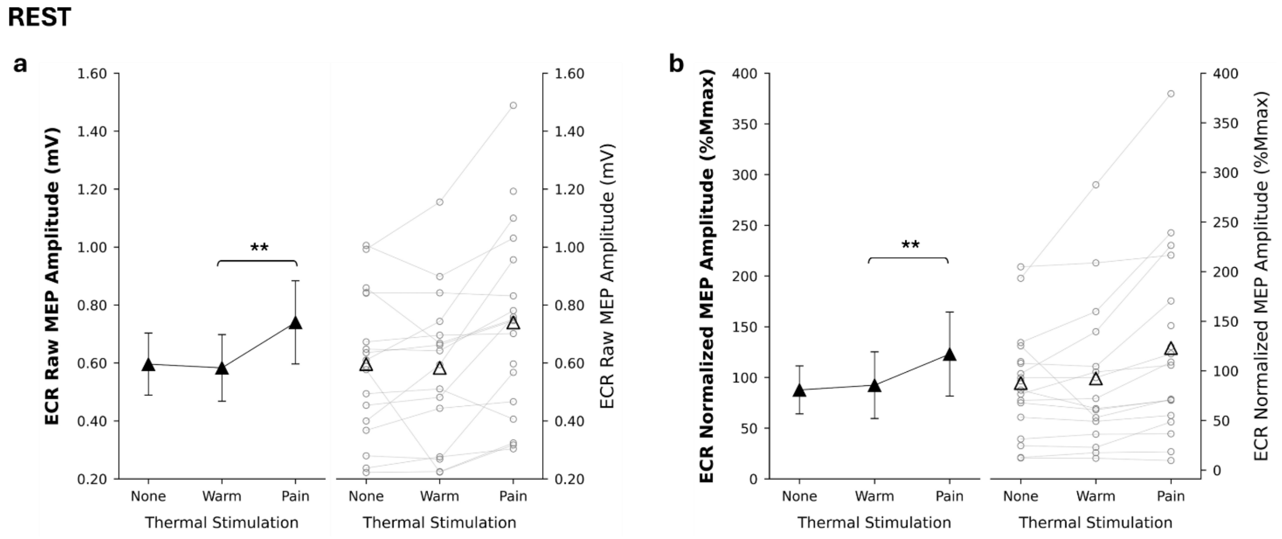


**Figure S 1: Effect of thermal stimulation on MEPs recorded from ECR muscle at rest.** In all panels, the left side shows mean values (black triangles) with 95% confidence interval error bars, while the right side shows individual data points (gray circles) along with the mean (black triangles). *Panel a* shows mean raw MEP. Pairwise comparisons revealed increased ECR raw MEP amplitude during pain compared to warm (p = .005). *Panel b* shows mean normalized MEP Pairwise comparisons revealed increased ECR normalized MEP amplitude during pain compared to warm (p = .009). ECR, extensor carpi radialis; MEP, motor evoked potential.

*ECR TS.* During rest, mean test stimulus amplitude was 0.84 ± 0.48 mV for control condition, 0.91 ± 0.44 mV for warm condition and 0.94 ± 0.44 mV for pain condition. No main effect of temperature was found on rest ECR test stimulus amplitude ($F$(1.97, 35.48) = 1.409, $p$ = .258, $\eta^2_p$ = .073 [.000, .236]).

*ECR SICI.* During rest, mean SICI as percentage of test stimulus was -38.34 ± 27.18 for control condition, -35.83 ± 22.25 for warm condition and -25.41 ± 27.36 for pain condition. Mean SICI as percentage of Mwave was -41.45 ± 45.95 for control condition, -52.85 ± 52.47 for warm condition and -42.29 ± 47.20 for pain condition. A main effect of temperature was found on SICI as percentage of the test stimulus ($F$(1.61, 29.06) = 3.617, $p$ = .048, $\eta^2_p$ = .167 [.000, .375]; **Fig. S2a**). Post hoc comparisons revealed reduced SICI during pain compared to warm conditions ($t$(18) = 2.841, $p$ = .033, $d$ = 0.406 [-0.011, 0.822]), with no differences between control and warm conditions ($t$(18) = -.454, $p$ = .655, $d$ = -0.098 [-0.667, 0.471]) and between control and pain conditions ($t$(18) = -2.217, $p$ = .079, $d$ = -0.503 [-1.142, 0.135]). No main effect of temperature was found on SICI as percentage of the Mwave ($F$(1.73, 31.06) = 0.950, $p$ = .386, $\eta^2_p$ = .050 [.000, .218]; **Fig. S2b**).

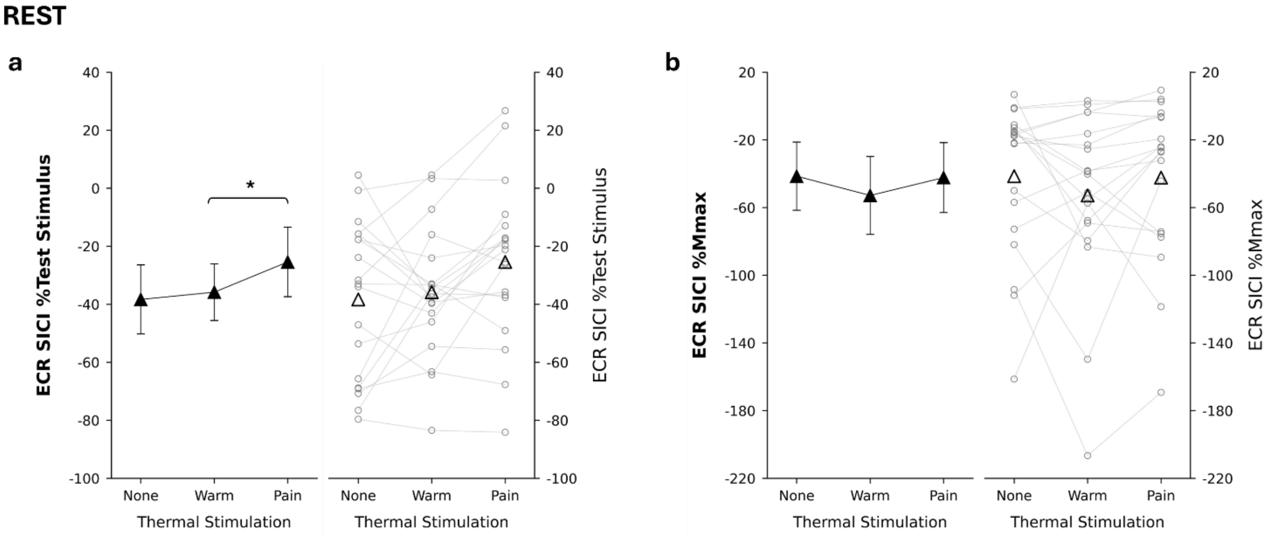


**Figure S 2: Effect of thermal stimulation on SICI recorded from ECR muscle at rest.** In all panels, the left side shows mean values (black triangles) with 95% confidence interval error bars, while the right side shows individual data points (gray circles) along with the mean (black triangles). *Panel a* shows mean SICI as %Test Stimulus. *Panel b* shows mean SICI as %Mwave. Repeated-measures ANOVAs revealed no main effect of thermal stimulation on SICI. ECR, extensor carpi radialis; Mwave, maximal compound action potential; SICI, short-interval intracortical inhibition.

*ECR ICF.* During rest, mean ICF as percentage of test stimulus was 54.97 ± 62.28 for control condition, 44.36 ± 44.62 for warm condition and 62.73 ± 46,60 for pain condition. Mean ICF as percentage of Mwave was 61.43 ± 88.86 for control condition, 54.56 ± 73.53 for warm condition and 83.52 ± 82.47 for pain condition. No main effect of temperature was found on ICF as percentage of the test stimulus ($F$(1.93, 34.66) = 1.211, $p$ = .309, $\eta^2_p$ = .063 [.000, .225]; **Fig. S3a**), nor on ICF as percentage of the Mwave ($F$(1.78, 32.10) = 3.094, $p$ = .064, $\eta^2_p$ = .147 [.000, .341]; **Fig. S3b**).

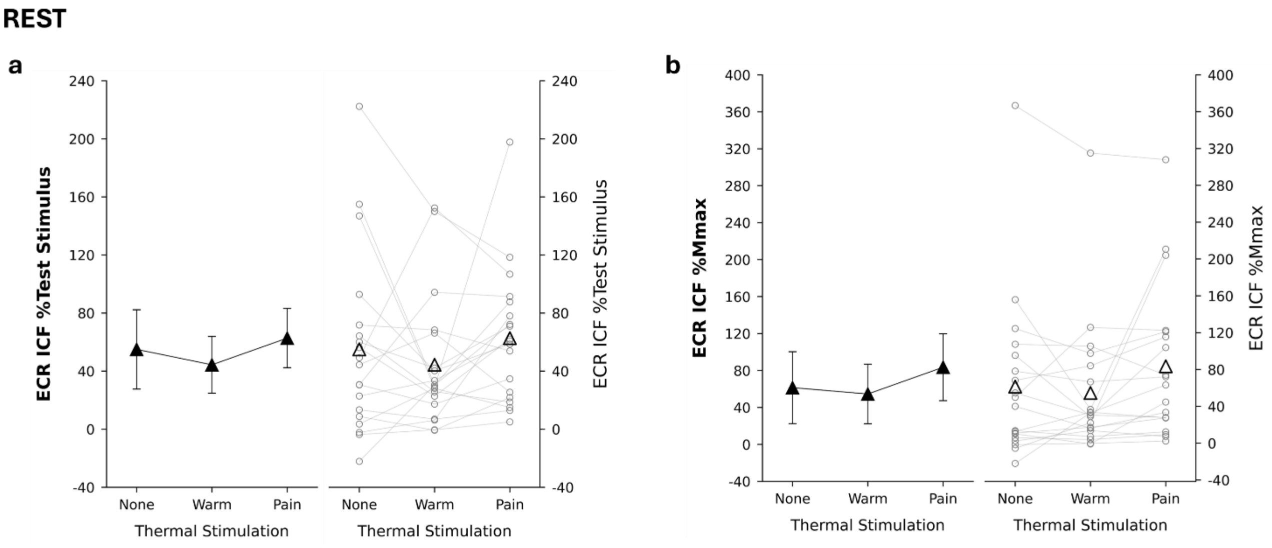


**Figure S 3: Effect of thermal stimulation on ICF recorded from ECR muscle at rest.** In all panels, the left side shows mean values (black triangles) with 95% confidence interval error bars, while the right side shows individual data points (gray circles) along with the mean (black triangles). *Panel a* shows mean ICF as %Test Stimulus. *Panel b* shows mean ICF as %Mwave. Repeated-measures ANOVAs revealed no main effect of thermal stimulation on ICF. ECR, extensor carpi radialis; ICF, intracortical facilitation; Mwave, maximal compound action potential.

### Supplementary material S3

#### Extensor carpi radialis (ECR) corticospinal and intracortical excitability during wrist flexions

*ECR MEP.* During voluntary wrist flexions, mean raw MEP amplitude was 1.49 ± 0.87 mV for control condition, 1.37 ± 0.65 mV for warm condition and 1.50 ± 0.68 mV for pain condition. Mean normalized MEP amplitude was 207.20 ± 143.30 for control condition, 190.90 ± 125.60 for warm condition and 203.1 ± 132.80 for pain condition. No main effect of temperature was found on raw MEP amplitude ($F(1.14, 22.88) = 1.932$, $p = .178$, $\eta^2_p = .088$ [.000, .322]; **Fig. S4a**), or on normalized MEPs amplitude ($F(1.21, 24.18) = 0.837$, $p = .391$, $\eta^2_p = .040$ [.000, .243]; **Fig. S4b**).

*ECR CSP.* During voluntary wrist flexions, mean raw CSP duration was 132.75 ± 35.51 ms for control condition, 139.85 ± 39.58 ms for warm condition and 136.35 ± 41.60 ms for pain condition. Mean normalized CSP duration was 143.90 ± 112.64 for control condition, 151.50 ± 137.64 for warm condition and 124.60 ± 82.32 for pain condition. No main effect of temperature was found on raw CSP duration ($F(1.12, 22.32) = 1.118$, $p = .309$, $\eta^2_p = .053$ [.000, .277]; **Fig. S4c**), or on normalized CSP duration ($F(1.59, 31.80) = 1.693$, $p = .204$, $\eta^2_p = .078$ [.000, .263]; **Fig. S4d**).

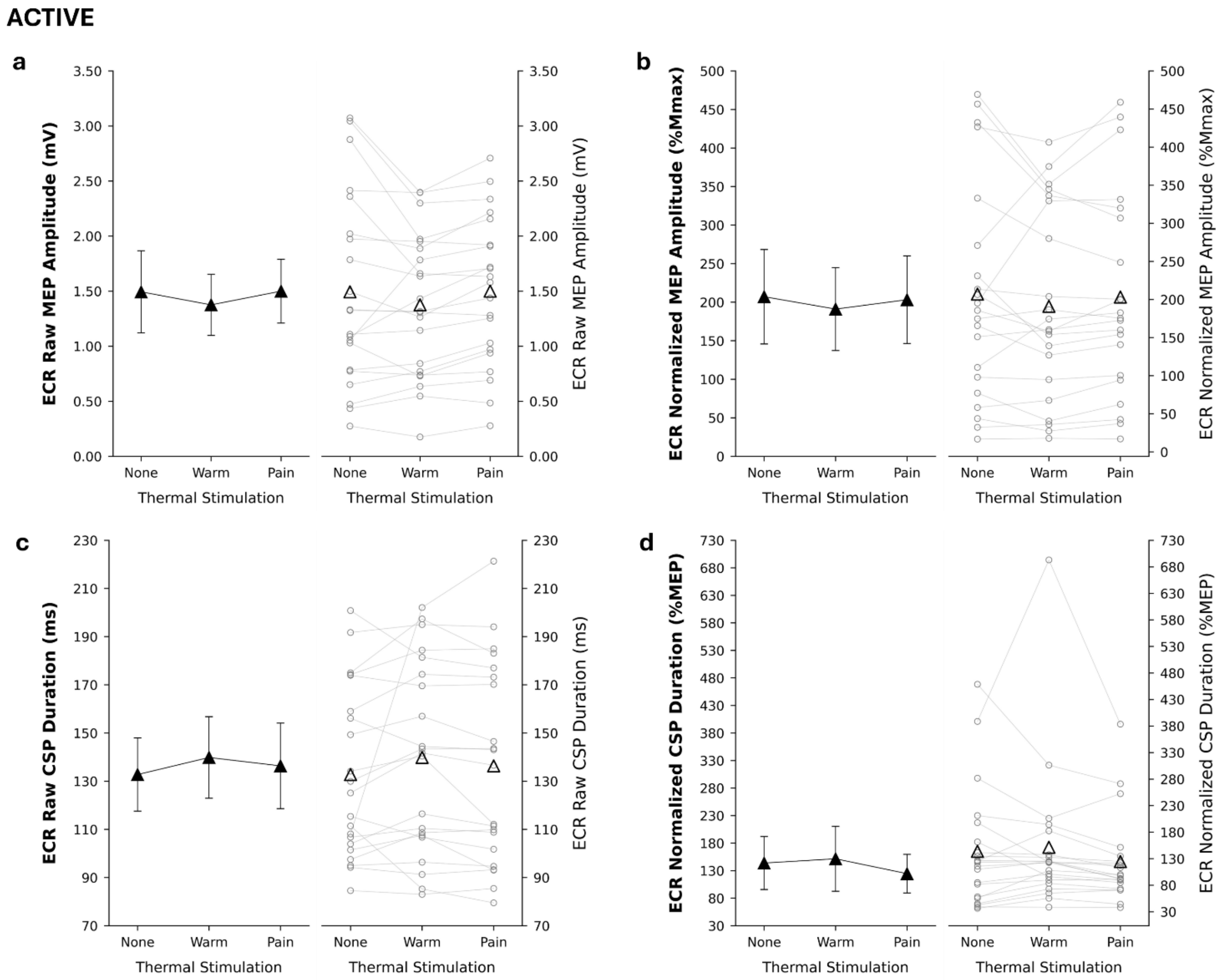


**Figure S 4: Effect of thermal stimulation on MEP and CSP recorded from ECR muscle during voluntary wrist flexions.** In all panels, the left side shows mean values (black triangles) with 95% confidence interval error bars, while the right side shows individual data points (gray circles) along with the mean (black triangles). *Panel a* shows raw MEP amplitudes. *Panel b* shows normalized MEP amplitudes. *Panel c* shows mean raw CSP duration. *Panel d* shows mean normalized CSP duration. Repeated-measures ANOVAs revealed no main effect of thermal stimulation on MEP and CSP. CSP, cortical silent period; ECR, extensor carpi radialis; MEP, motor evoked potential.

*ECR TS.* During active condition, mean test stimulus amplitude was 1.25 ± 0.69 mV for control condition, 1.23 ± 0.84 mV for warm condition and 1.22 ± 0.69 mV for pain condition. No main effect of temperature was observed on test stimulus amplitude ($F(1.51, 30.18) = 0.137$, $p = .814$, $\eta^2_p = .007$ [.000, .115]).

*ECR SICI.* During voluntary wrist flexions, mean SICI as percentage of test stimulus was -7.29 ± 31.85 for control condition, -1.96 ± 51.70 for warm condition and -1.38 ± 27.74 for pain condition. Mean SICI as percentage of Mwave was -38.76 ± 87.78 for control condition, -26.47 ± 72.47 for warm condition and -18.52 ± 63.02 for pain condition. No main effect of temperature was found on SICI expressed as a percentage of test stimulus ($F(1.67, 33.44) = 0.582$, $p = .535$, $\eta^2_p = .028$ [.000, .172]; **Fig. S5a**) or as a percentage of Mwave ($F(1.95, 39.01) = 1.397$, $p = .259$, $\eta^2_p = .065$ [.000, .264]; **Fig. S5b**).

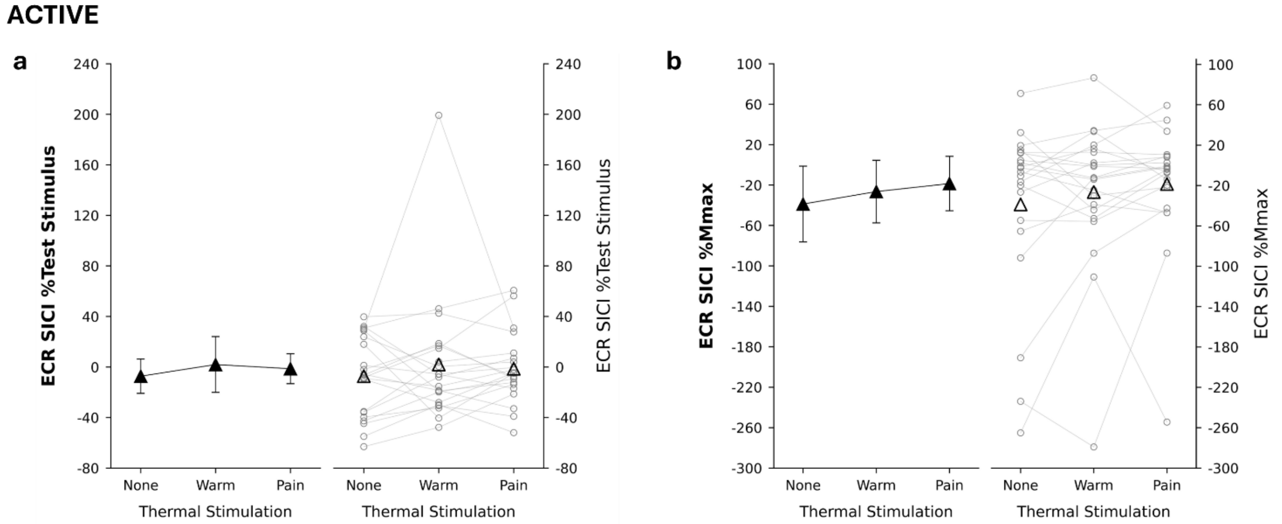


**Figure S 5: Effect of thermal stimulation on SICI recorded from ECR muscle during voluntary wrist flexions.** In all panels, the left side shows mean values (black triangles) with 95% confidence interval error bars, while the right side shows individual data points (gray circles) along with the mean (black triangles). *Panel a* shows mean SICI as %Test Stimulus. *Panel b* shows mean SICI as %Mwave. Repeated-measures ANOVAs revealed no main effect of thermal stimulation on SICI. ECR, extensor carpi radialis; Mwave, maximal compound muscle action potentials; SICI, short-interval intracortical inhibition.

*ECR ICF.* During voluntary wrist flexions, mean ICF as percentage of test stimulus was 29.12 ± 35.84 for control condition, 23.45 ± 41.09 for warm condition and 32.61 ± 52.72 for pain condition. Mean ICF as percentage of Mwave was 50.92 ± 61.53 for control condition, 45.15 ± 77.80 for warm condition and 65.44 ± 89.88 for pain condition. No main effect of temperature was found on ICF expressed as percentage of test stimulus ($F(1.53, 30.66) = 0.531$, $p = .547$, $\eta^2_p = .026$ [.000, .179]; **Fig. S6a**) or as a percentage of Mwave ($F(1.98, 37.58) = 0.942$, $p = .398$, $\eta^2_p = .047$ [.000, .192]; **Fig. S6b**).

**ACTIVE**

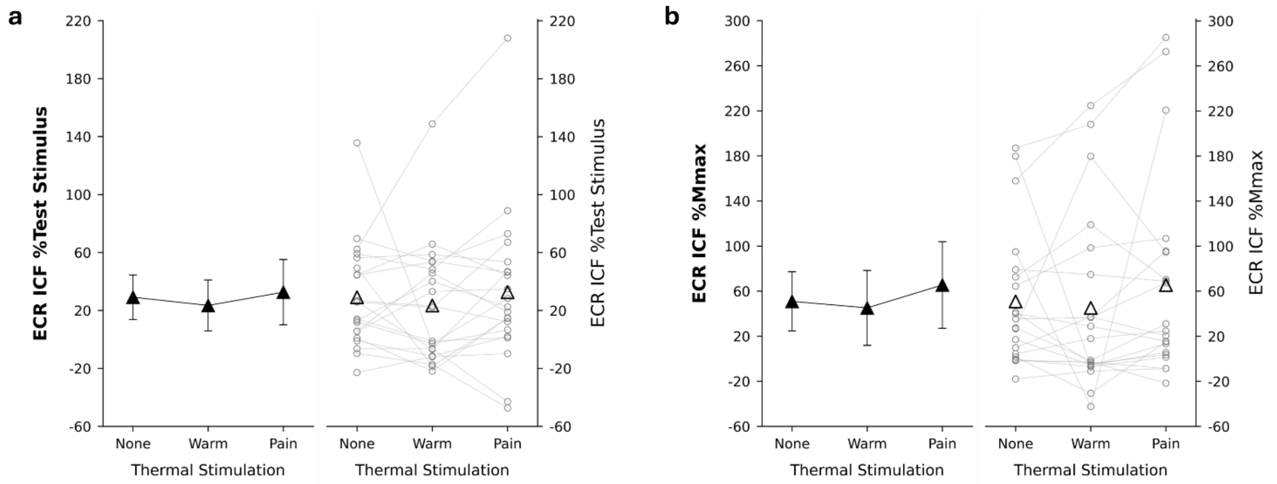


**Figure S 6: Effect of thermal stimulation on ICF recorded from ECR muscle during voluntary wrist flexions**. In all panels, the left side shows mean values (black triangles) with 95% confidence interval error bars, while the right side shows individual data points (gray circles) along with the mean (black triangles). *Panel a* shows mean ICF as %Test Stimulus. *Panel b* shows mean ICF as %Mwave. Repeated-measures ANOVAs revealed no main effect of thermal stimulation on ICF. ECR, extensor carpi radialis; ICF, intracortical facilitation; Mwave, maximal compound muscle action potentials.
